# Multisensory integration for active mechanosensation in *Drosophila* flight

**DOI:** 10.1101/2025.06.20.660728

**Authors:** Kevin M. Mills, Noah J. Cowan, Marie P. Suver

## Abstract

To support robust behaviors in highly variable environments, animals rely on active sampling of their sensory surroundings. Here, we use tethered, flying *Drosophila melanogaster* and a multisensory behavioral apparatus simulating forward flight to determine how visual and mechanosensory information are integrated and control active movements of an important multimodal sensory organ, the antennae. We found that flies perform active antennal movements in response to varying airflow, and that the direction of these movements changes depending on the visual environment. Next, we found that antennal movements are amplified in the presence of visual motion, but only when the fly was flying. Through mechanical and optogenetic manipulation of mechanosensory input, we found that mechanosensory feedback is vital to antennal positioning at flight onset. Additionally, we observed unexpected changes in wingbeat frequency when the antenna was mechanically stabilized, suggesting that multiple antennal mechanosensors contribute to flight regulation. Finally, we show that integration of mechanosensory and visual cues for controlling antennal motion follows in a “winner-takes-all” paradigm dependent on the stimulus frequency, mirroring visuo-mechanosensory guided behaviors in other species. Together, these results reveal novel behavioral gating of sensory information and expand our understanding of the efferent control of active sensing.

## INTRODUCTION

An important function of nervous systems is the ability to quickly and efficiently integrate sensory information from multiple sensory modalities. Although a single sensory modality can be sufficient to guide motor actions in isolation (e.g. bacterial chemotaxis^1^), complex animal behavior typically relies on several sensory modalities whose reliability varies depending on environmental conditions. Many behaviors rely on multisensory integration, such as host-seeking^2^, the righting reflex (e.g. in hoverflies)^3^, and perception of self-motion^4^. To reconcile variably useful sensory information from the environment, nervous systems implement different integrative mechanisms. In some cases, this integration is approximately the linear superposition of signals from distinct modalities^5,6^ (but see Oie et al.^7^). However, in other instances, one sensory modality dominates (i.e. ‘winner-takes-all’) in its influence on motor commands^8,9^.

Beyond integrating input from multiple sensory modalities, animals dynamically alter how they interact with and interpret their sensory environment through active movements^10^. These ‘active sensing’ behaviors, in which animals use muscles to position their sensors, directly modify how sensory information is acquired and can aid in effective sensation for guiding behavior^11^. For example, rodents perform rhythmic, low-frequency movements of their whiskers to sense objects in their environment during goal-directed navigation. This active sampling of information (‘whisking’) influences incoming sensory input through sensory filtering^12^ (‘gating’) and guides future sampling movements^13^. Active sensing behaviors are abundant across phyla^14–18^, but insects employ a remarkably conspicuous and diverse array of active antennal behaviors to sense their environment. For instance, cockroaches repeatedly touch and investigate objects of interest with their antennae^19^, locusts alter antennal movements to improve odor encoding for olfactory-guided behavior^20^, and several insects use antennal contact-chemosensation in behaviors such as courtship^21^ and odor trail navigation^22^.

Insect antennae are highly multimodal, and house many sensors supporting olfaction^23^, hygrosensation and thermosensation^24,25^, and mechanosensation^26–29^. Thus, active antennal movements can influence multiple sensory modalities. These active antennal movements are guided by musculature located at antennal joints that enable directed acquisition of sensory information^30–36^. This recurrent interaction between antennal motion and antennal sensation is critical for many insect behaviors, including tactile exploration^37,38^, escape^39^, oviposition^40^, and flight^41–43^. In addition to housing multiple senses, antennal movements are likely driven by multimodal information. Yet how different sensory inputs guide the movements of this dynamic sensor are not well understood.

In the fruit fly *Drosophila melanogaster*, antennal mechanosensation is crucial for flight control and navigation. A large population of mechanosensory stretch-receptive neurons, called Johnston’s Organ Neurons^44^ (JONs), detect passive motion of the antennae induced by both high-frequency air vibrations (e.g. courtship song^45,46^) and static airflow^47,48^. Flies use this mechanosensory information to guide several flight behaviors including orientation^49^, long-term navigation^50^, and groundspeed regulation^51^. Flight is also heavily reliant on vision, which is crucial for stabilizing reflexes^52^, rapid turns known as saccades^53–56^, altitude control^57^, obstacle avoidance^58^, and groundspeed regulation^51,59^. Further, evidence from previous studies demonstrated that antennal movements are regulated by both mechanosensory^35^ and wide-field visual^60^ sensory input. However, it is not fully understood how the brain integrates these two – often complementary – sensory modalities to regulate antennal positioning for important active sensing behaviors.

In this study, we used a custom multisensory behavioral apparatus to determine the role of mechanosensory and visual information in guiding antennal movements and forward flight regulation in tethered flying and nonflying fruit flies. We found that antennal position is actively regulated in response to varying airspeeds regardless of whether the animal was flying or not. We also measured the influence of mechanosensory input on airflow-dependent antennal positioning and wingbeat kinematics by blocking antennal mechanosensory input and found that blocking antennal mechanosensation altered flight-dependent antennal positioning and wingbeat frequency. Additionally, we assessed the influence of progressive visual motion on active antennal movements and found that these responses are gated by flight state. Finally, we presented sinusoidal patterns of frontal airflow and progressive optic flow to gain a deeper understanding of how these multisensory stimuli are integrated by the antennal sensorimotor system and used this to build a model of the neural computations contributing to this integration.

## RESULTS

### Flies actively position their antennae in response to frontal airflow

To assess how mechanosensory and visual stimuli influence antennal movements and wingbeat kinematics during forward flight, we built a behavioral apparatus that delivers controlled frontal airflow and wide-field progressive optic flow along the anterior-posterior axis. We rigidly mounted flies to thin tungsten pins, fixed the head in its resting position, and placed them in the center of the behavioral apparatus. To record wingbeat information, we mounted a light-based wingbeat sensor posterior to the fly and recorded antennal movements using a camera positioned dorsal to the fly (Figure 1A, STAR Methods).

**Figure 1.**
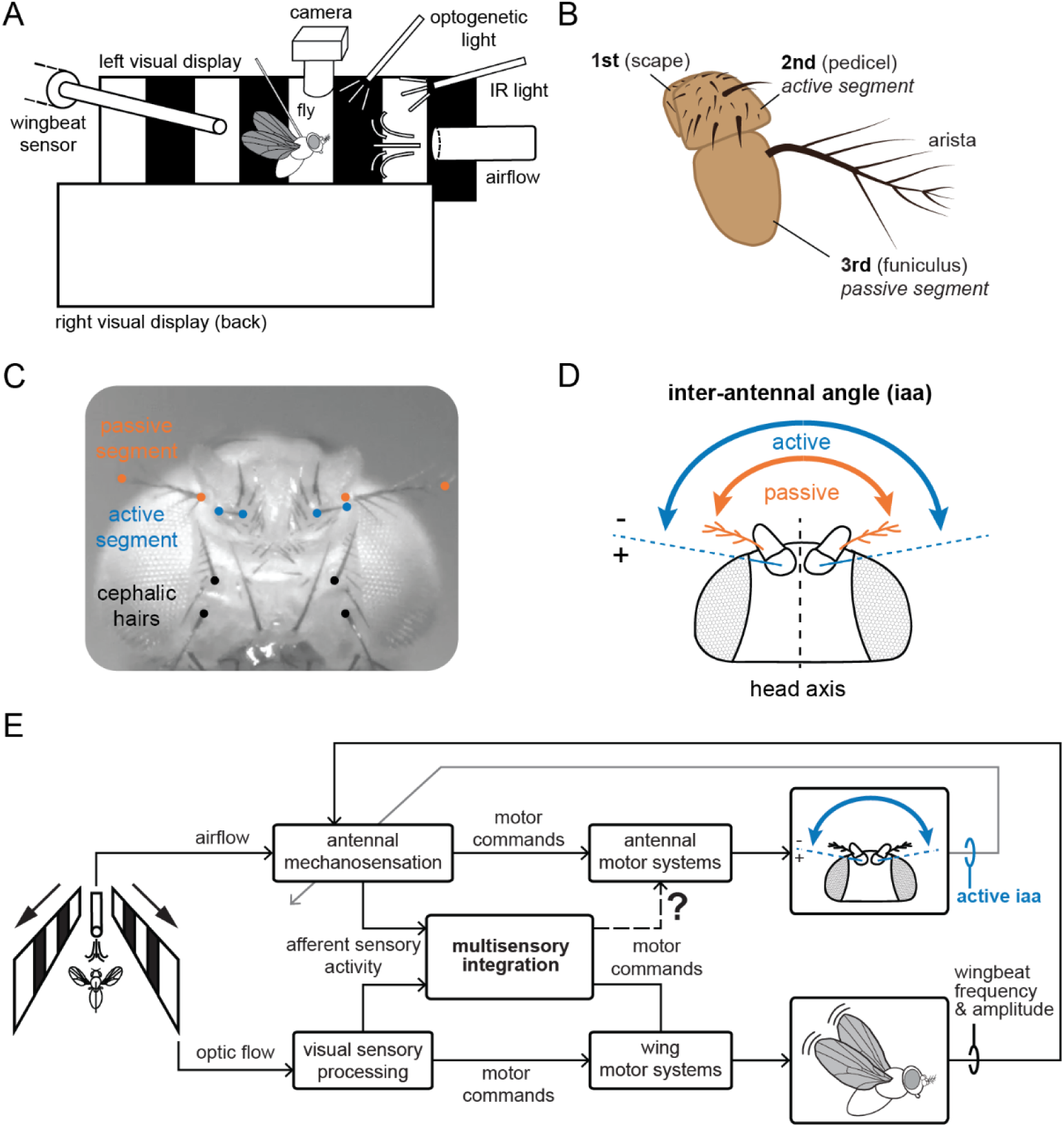
Multisensory behavioral apparatus and experimental design. A) Schematic of the behavioral apparatus (not to scale). Fly is rigidly tethered to a pin and placed directly in the path of a frontal airflow tube. Visual displays are equidistant from the fly on the left and right side. Infrared light illuminates the fly from the front, and a wingbeat sensor detects light reflected by the wings from behind. A camera captures antennal movements from above the fly (dorsally). B) Schematic of the fly antenna, showing the three primary segments: first (scape), second (pedicel), and third (funiculus). Additionally, the arista (rigidly attached to third segment) is shown. C) Image of the fly indicating points tracked on both the head and antennae, including a hair on the second segment (blue), the arista (orange), and the base of four cephalic hairs (black). D) Schematic indicating measure of inter-antennal angle (‘iaa’) for the passive (orange) and active (blue) segments of the antennae. E) Block diagram depicting our experimental approach. The bottom branch illustrates the visuomotor transformations that drive changes in wing kinematics (e.g., wingbeat frequency and amplitude) during tethered flight^51^. Wing vibrations are detected by the antennae, illustrated by the feedback from the wings to the antennal mechanosensation block. Additionally, antennal movements may modulate wind sensing, as illustrated by the “active iaa” feedback to the mechanosensory block. This paper characterizes the mechanosensory control of the antennal motor control loop, illustrated by the dashed branch (labeled by the ‘?’).

We tracked several points on the second and third antennal segments (also referred to as ‘pedicel’ and ‘funiculus’, respectively) to quantify antennal movements (Figure 1B, DeepLabCut^61^). To measure passive motion, we tracked the base and tip of the arista (Figure 1C and 1D, orange) a large, rigidly fixed hair-like appendage attached to the third segment that transduces incoming airflow, amplifying passive motion much like a sail^28,62,63^. To measure active motion, we tracked the tip and base of a large hair (Figure 1C and 1D, blue) on the second segment, which is actuated by four muscles located in the first segment^34,35^ (Figure 1B, also called the ‘scape’). We measured the angles of both the active and passive segments relative to the midline of the head. Because both the visual and mechanosensory stimuli were bilaterally symmetric, we added the angle of the left and right antennae relative to the midline together to compute the inter-antennal angle (‘iaa’, Figure 1D), a measure that has been extensively used in the past for gauging antennal position across insect species^32,42,64,65^, for both the passive and active segments.

A block diagram (Figure 1E) illustrates our experimental approach, the sensory stimuli we presented, and behavioral outputs we measured. Because the fly is tethered and the stimuli are presented in open-loop, changes to wingbeat kinematics do not drive reafferent visual feedback^66^. However, antennal movements generate two forms of reafferent feedback: (1) the motor commands to the antenna drive changes to inter-antennal angle, potentially modulating wind sensing (Figure 1E, ‘active iaa’ feedback to mechanosensory block), and (2) changes in wingbeat kinematics can be detected by the antennae through high frequency vibrations^47,67^ (Figure 1E, wing feedback to mechanosensory block).

To evaluate how flies actively position their antennae in varying airflow, we measured both passive and active segment inter-antennal angles in response to wind speeds ranging from 0-300 cm/s. We presented all wind speeds in either ascending or descending order for 6 sec, for a total of 10 trials per fly (pseudo-randomized order, see STAR Methods). To confirm that the active antennal segment movements we measured were not a product of passive displacement by wind, we performed the same experiment with freshly dead flies. In both dead flies and live flies in darkness (i.e., visual stimulus off), we observed an increase in the inter-antennal angle of the passive segment with increasing wind speed (Figure 2A). Whereas the inter-antennal angle for the active segment reached a maximal increase of 2.72 deg at the highest wind speeds in dead flies (Figure 2B, light gray trace), live nonflying flies in darkness exhibited the opposite response, decreasing their inter-antennal angle as wind speeds increased (Figure 2B, dark gray trace). This observation suggests that the active antennal segment was passively displaced at most ∼3 deg apart at higher wind speeds. In live flies, however, active movements typically exceed the passive deflection that would be created by aerodynamic forces, and these movements vary as a function of wind speed.

**Figure 2.**
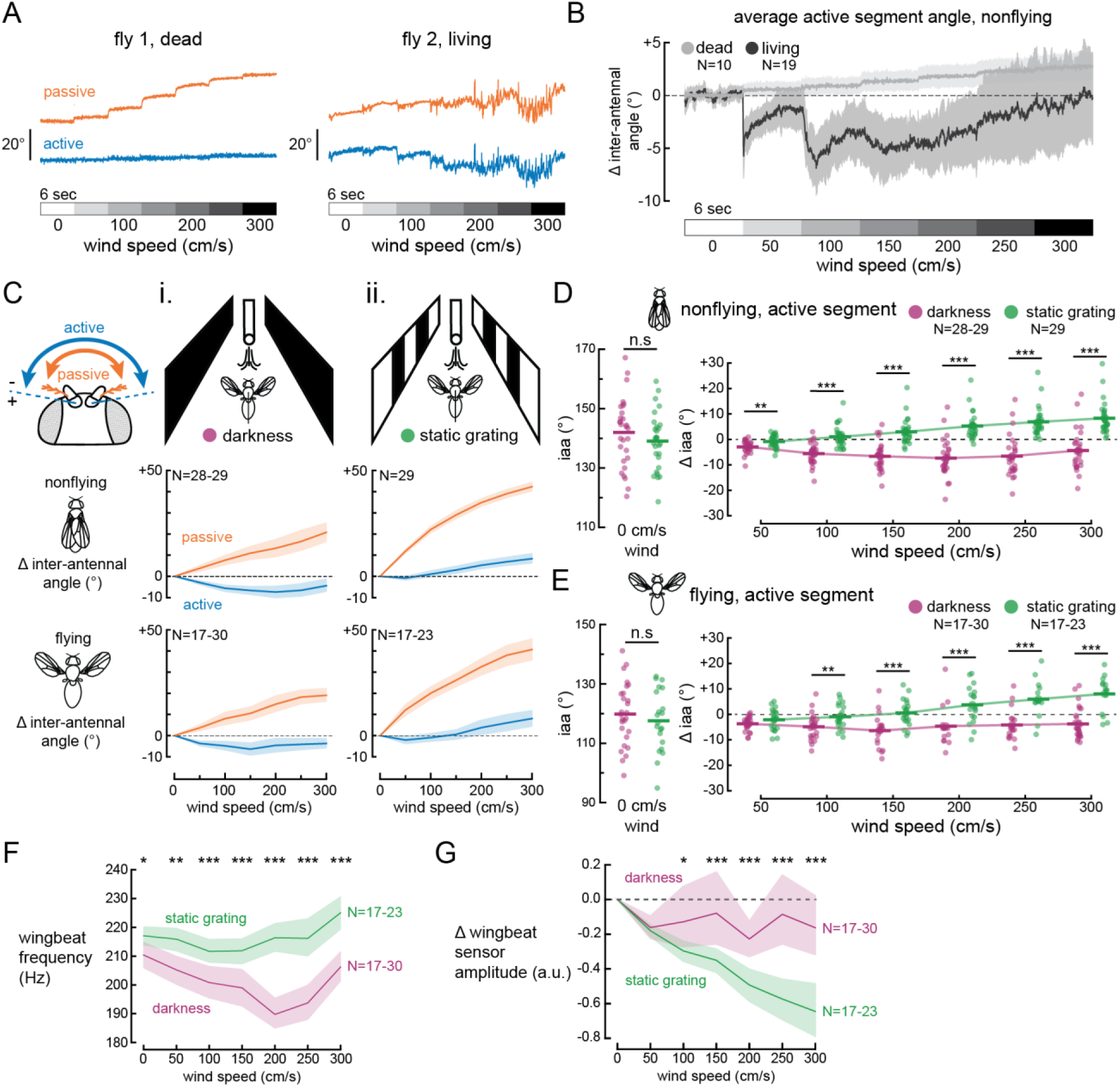
Airflow-dependent active positioning of the antennae is reversed in the presence of a static grating. A) Example single-trial traces of passive (orange) and active (blue) antennal movements for a dead fly (left) and a live fly (right) in darkness; gradient bar indicates wind speed increases every six sec. B) Average active antennal positioning over time as wind speed increases for dead (light grey) and live, nonflying flies (dark grey). All movements are baseline-subtracted relative to the average angle at 0 cm/s wind speed. C) Average passive and active inter-antennal angles, baseline-subtracted from 0 cm/s wind speed, for flies in darkness and with a static grating. D) Active inter-antennal angle across wind speeds for nonflying flies in darkness (purple) and with a static grating (green). Left: absolute inter-antennal angle at 0 cm/s wind speed (p=.30). Right: baseline-subtracted (i.e., relative to 0 cm/s wind speed) inter-antennal angle across wind speeds in both conditions. Flies in darkness moved their antennae inwards as wind speed increased (β=-.016, p<.001), while flies with a static grating moved theirs outward (β=.048, p<.001). E) Similar to (D), for flying flies. As wind speed increased, flies in darkness positioned their antennae inwards (β=-.010, p<.01), while those with a static grating positioned theirs outward (β=.030, p<.001). F) Average wingbeat frequency for both conditions. Flies with a static grating had a higher wingbeat frequency across wind speeds (all p<.05). G) Average relative wingbeat amplitude decreased as wind speed increased (see Figure S1). Wingbeat amplitude decreased more for flies with a static grating (β=-1.97e-3, p<.001) and was significantly lower at and above 100 cm/s wind speed (all p<.05).

### Airflow-dependent active antennal positioning response reverses in the presence of a static grating

To determine how the visual environment influences wind-induced active movements, we presented flies with a static grating of black and white bars equidistant on their left and right (Figures 1A and 2C). In darkness, we observed that both nonflying and flying flies decreased their active inter-antennal angle as wind speeds increased (Figure 2Ci, blue traces). However, in the presence of a static grating, this trend reversed; active segment inter-antennal angle decreased slightly only at the lowest tested airflow (50 cm/s) before increasing with higher wind speeds (Figure 2Cii, blue traces). Furthermore, this alteration of wind speed-dependent active positioning directly affected the positioning of the passive segment; flies in the presence of a static grating exhibited greater passive inter-antennal angle increases with higher airflow compared to flies in darkness (Figure 2C, orange traces).

In the absence of wind, we observed a small but statistically nonsignificant decrease in absolute active antennal position for flies in darkness compared to those presented with a static grating; this was true for both nonflying (Figure 2D, left; p=.30, t-test) and flying flies (Figure 2E, left; p=.46, t-test). Upon exposure to airflow, nonflying flies in darkness positioned their antennae inwards (Figure 2D, right; p<.001, linear-mixed model). This trend reversed in flies presented with a static grating, which positioned their antennae outward with increasing wind speed (p<.001). Moreover, flies in the presence of a static grating positioned their antennae outward with increased wind speed at approximately twice the rate (+1.6 deg per 50 cm/s wind speed) flies in darkness positioned their antennae inwards (-0.8 deg per 50 cm/s). Post-hoc comparisons revealed significant differences in relative inter-antennal angle between the two visual conditions at all wind speeds above 0 cm/s (all p<.01, Holm-corrected). In flying flies, we observed a similar trend in antennal positioning as in nonflying flies: flies in darkness positioned their antennae inwards as wind speeds increased (p<.01, linear-mixed model) whereas flies with a static grating positioned their antennae outward (p<.001). These differences in relative inter-antennal angle between the two visual conditions were statistically significant at and above 100 cm/s wind speed (all p<.01, Holm-corrected).

### Airflow-dependent wingbeat dynamics are modulated by the visual environment

Previous studies demonstrated that active antennal movements occur during flight^35,60^, so we next asked how wing dynamics might change in response to the same stimuli that elicit airflow-dependent antennal movements. Overall, we found that flying flies exhibited a variety of wing dynamics dependent on both wind speed and the visual environment. We observed that flies in darkness decreased their wingbeat frequency as wind speed increased up to 200 cm/s (Figure 2F; p<.001, linear-mixed model). However, flies in the presence of a static grating exhibited a subtle increase in wingbeat frequency with higher wind speeds (p<.001 compared to flies in darkness, linear-mixed model). Additionally, flies presented with a static grating had a higher wingbeat frequency than those in darkness across all wind speeds (all p<.01, Holm-corrected). We also observed a decrease in relative wingbeat amplitude (Figure S1, STAR Methods) as wind speed increased in both visual conditions (in darkness; p=.006, linear-mixed model), although flies in the presence of a static grating exhibited a larger, more consistent negative trend (p<.001). There was a significant difference in relative wingbeat amplitude between visual conditions at and above 100 cm/s wind speed (all p<.05, Holm-corrected). We confirmed our measure of wingbeat amplitude in a separate experiment (Figure S1, STAR Methods).

### Relative active antennal positioning across wind speeds is flight-state invariant

Flying insects actively position their antennae inwards during flight^26,65,68–70^. To determine whether airflow-dependent active antennal movements in *Drosophila* are altered during flight, we compared the responses of nonflying and flying flies to varying wind speed both in darkness and the presence of a static grating (Figure 3). To quantify this movement, we measured the average position of the aristae in nonflying and flying flies (Figure 3A and 3B). We found that the inter-antennal angle of the arista was 121.7 ± 5.1 deg and 97.9 ± 6.5 deg (mean ± SD) in nonflying and flying flies, respectively. Because we observed no significant differences in inter-antennal angle between visual conditions in both flight states without airflow (Figure 2D-E, left), we combined data from both visual conditions for this analysis.

**Figure 3.**
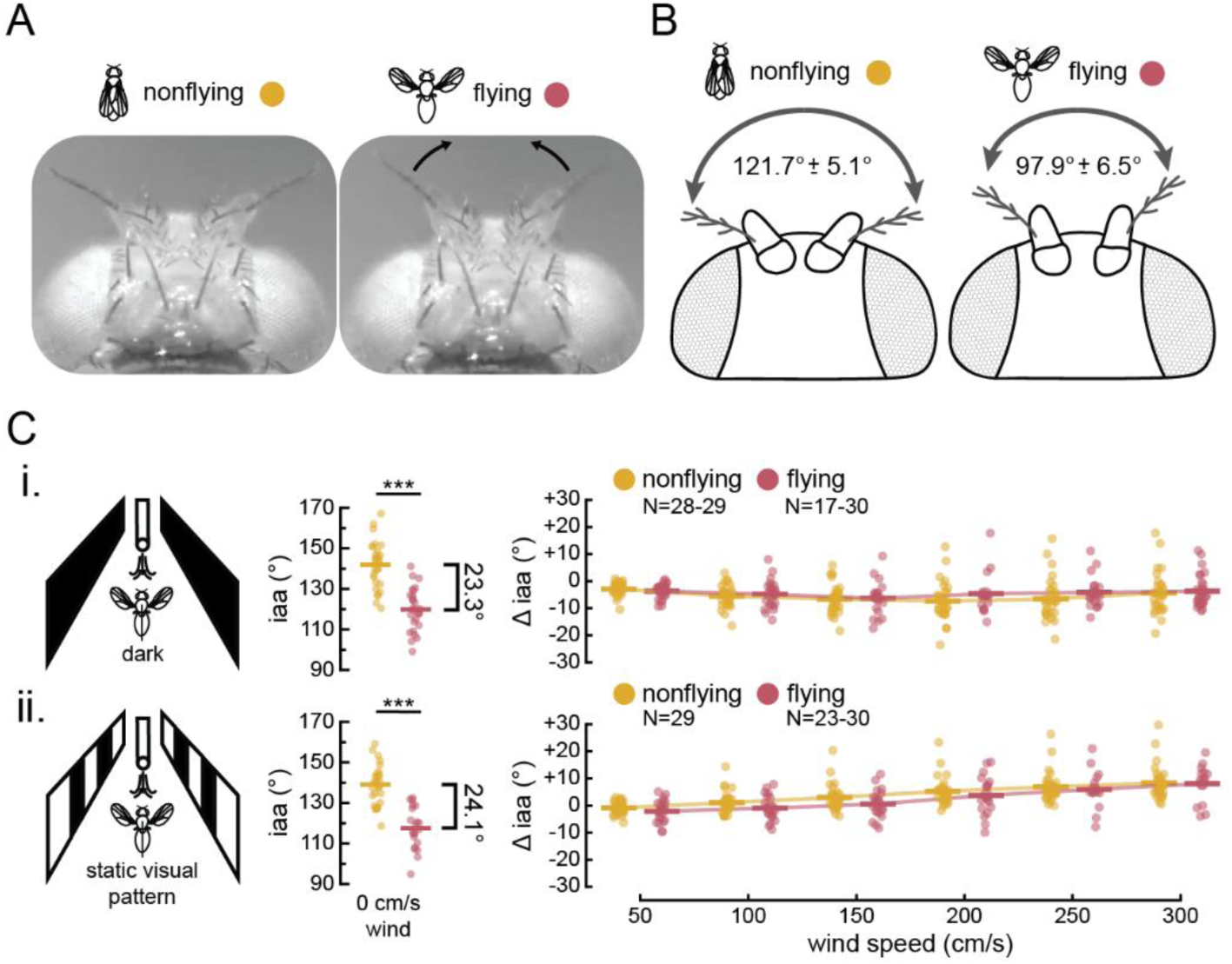
Active frontal antennal positioning at the onset of flight is independent of airflow-dependent active movements. A) Dorsal view of the fly’s head during nonflight (left) and flight (right), showing frontal antennal movement at flight onset. B) Schematic of frontal positioning from (A), with average inter-antennal angle ± standard deviation for nonflying (left) and flying (right) flies (N=52 flies). C, i) Active antennal positioning across wind speeds for flies in darkness. Left shows absolute active inter-antennal angle for nonflying (yellow) and flying (red) flies (mean difference = 23.3 deg, p<.001). Right shows baseline-subtracted (i.e., relative to 0 cm/s wind speed) active inter-antennal angle across wind speeds. Wind speed-dependent antennal responses did not change in flight, despite inwards positioning at flight onset (β=-6.36e-3, p=.18). C, ii) Similar to (C, i), but flies are in the presence of a static grating. Left shows absolute active inter-antennal angles (mean difference = 24.1 deg, p<.001). Wind speed-dependent antennal responses did not change in flight (β=-1.97e-3, p=.60).

In the absence of wind, active segment inter-antennal angle was significantly lower during flight in both darkness and with a static grating (Figure 3C, middle; both p<.001, paired t-tests). However, flight state did not affect relative active antennal positioning across wind speeds in darkness (Figure 3Ci, right; p=.18, linear-mixed model) or in the presence of a static grating (Figure 3Cii, right; p=.60). Furthermore, we observed no significant differences in relative inter-antennal angles between nonflying and flying flies across all wind speeds for both visual conditions (all p>.05, Holm-corrected). These findings suggest that airflow-dependent active movements of the antennae are not substantially altered during flight.

### Silencing antennal sensory neurons shifts active antennal positioning more frontally at flight onset

Our findings thus far indicated that mechanosensory input influences active positioning of the antennae and wingbeat dynamics. To better understand the sensory influence on antennal movements, we measured antennal positioning responses to airflow in flies with reduced sensory input from the antennae by expressing GtACR1^71^, an inhibitory channelrhodopsin, in all Johnston’s Organ Neurons (JONs; *nan-GAL4>UAS-GtACR1*)^72,73^ (Figure 4A, yellow, ‘inactivated’). As an alternate method of abolishing JON activity, we glued the active-passive joint of both antennae (Figure 4A, blue, ‘glued’). This manipulation prevents the deflection of the passive segment in response to wind, effectively silencing JON activity^67^. To control for the presence of cyan light in our optogenetic inactivation experiments, we presented intact and glued flies with the same light stimulus used to invoke optogenetic silencing in the inactivated group.

**Figure 4.**
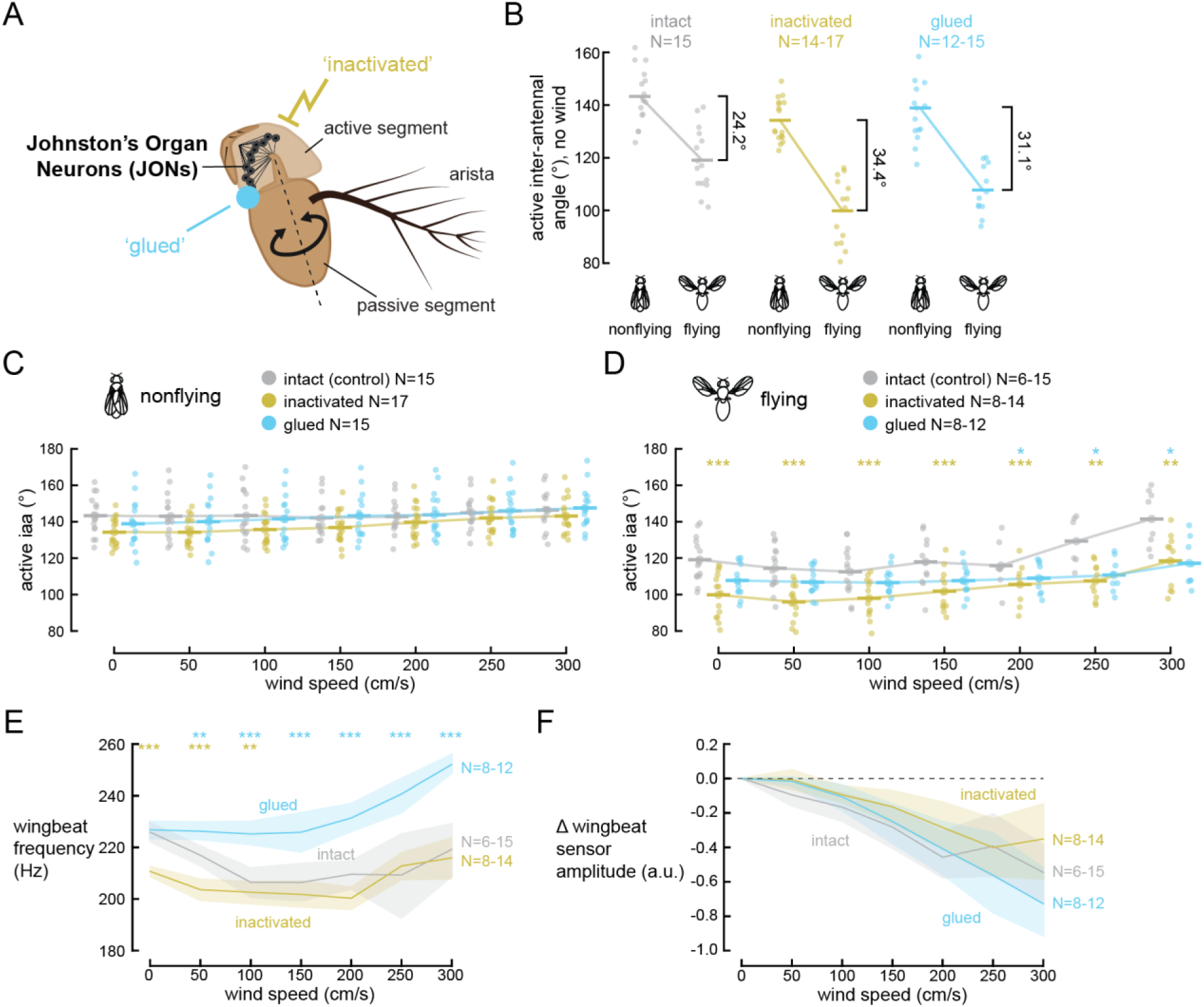
Blocking antennal sensations disrupts active frontal positioning of the antennae. A) Schematic of the *D. melanogaster* antenna highlighting the active segment (pedicel), passive segment (funiculus), and arista. The dashed line indicates the rotation axis of the passive segment, which activates stretch receptive Johnston’s Organ Neurons (JONs). JON activity is silenced optogenetically (*nan-GAL4>UAS-GtACR1*, yellow) or mechanically by gluing the active-passive segment joint (blue). B) Absolute active segment inter-antennal angle in flies with intact JONs (grey), optogenetically inactivated JONs (yellow), and mechanically silenced JONs (blue) in no airflow. Optogenetic inactivation resulted in a significantly larger inwards antennal movement (β=-9.26, p<.001) while gluing did not (β=-2.06, p=.40). C) Absolute active inter-antennal angle across all wind speeds in nonflying flies. No differences were found between conditions at all wind speeds (all p>.05). D) Same as (C), but in flying flies. Optogenetic inactivation resulted in significantly more inwards antennal positioning across all wind speeds (all p<.01). Gluing the active-passive joint resulted in more inwards antennae above 200 cm/s wind speed (all p<.05). E) Wingbeat frequency varies across conditions and wind speeds. Notably, wingbeat frequency decreases below 100 cm/s in flies with inactivated JONs (all p<.05) and increases above 50 cm/s when the active-passive joint is glued (all p<.01). F) Wingbeat amplitude decreases with wind speed in all conditions. Glued antennal joints result in a steeper decline (β=-6.61e-4, p=.012), but post-hoc comparisons revealed no differences across conditions (all p>.05).

We first investigated active movements across conditions in the absence of airflow to determine if JON silencing affects the active frontal antennal movement at flight onset. In the light control group (*Canton-S>UAS-GtACR1*) with no frontal airflow, we observed inwards positioning of the antennae (-24.2 deg, p<.001, linear-mixed model) at flight onset (Figure 4B, grey, ‘intact), similar in magnitude to movements observed in wild-type flies (-23.3 deg and -24.1 deg, Figure 3C, middle). However, in flies with optogenetically inactivated JONs, we observed significantly greater inwards positioning of the antennae (-34.4 deg, Figure 4B, yellow; p<.001, linear-mixed model) during flight. Likewise, we measured a similar increase in inwards antennal positioning in flies with glued active passive joints (-31.1 deg, Figure 4B, blue), although this relationship was not significant when compared to intact flies (p=.40, linear-mixed model).

The above findings demonstrate that in the absence of wind, blocking antennal sensation alters the inward positioning of the antennae at the onset of flight. To understand how reduced mechanosensation affected antennal movements in the presence of airflow stimuli, we next investigated active antennal positioning in response to a range of wind speeds with silenced JONs. In nonflying flies, we observed an increase in active inter-antennal angle as wind speed increased in all conditions (Figure 4C; all p<.001, linear-mixed model). This wind speed-dependent trend had a significantly higher slope in flies with optogenetically (+1.64 deg per 50 cm/s wind speed) and mechanically (+1.42 deg per 50 cm/s) silenced JONs relative to flies with intact JONs (+0.44 deg per 50 cm/s). However, we did not observe any significant differences in absolute antennal position between flies with intact JONs and flies in either silencing condition at any single wind speed (all p>.05, Holm-corrected).

In flying flies, active inter-antennal angle also increased at higher wind speeds in all conditions (Figure 4D). However, flies with glued active-passive joints displayed slower outward movement with increasing speed compared to flies with intact and inactivated JONs (p=.023, linear-mixed model). Notably, we observed further inwards positioning of the antennae when JONs were silenced (Figure 4D), similar to the response in the absence of airflow (Figure 4B, blue). Flies exhibited this narrower inter antennal angle all wind speeds for flies with optogenetically inactivated JONs (all p<.01, Holm-corrected) and at higher wind speeds for flies with mechanically silenced JONs (all p<.05). Together, these results suggest that JON sensation contributes to the antennal set position during flight, independent of wind speed.

### Complex effects of antennal mechanosensation manipulation on flight control

To determine how altered antennal mechanosensation affected flight control, we investigated wing dynamics across wind speeds in flies with silenced JONs. We found that flies with glued active-passive antennal joints have a higher wingbeat frequency than control flies at all wind speeds at and above 50 cm/s (Figure 4E, blue; all p<.001, Holm-corrected). Unexpectedly, we did not observe this trend in flies with optogenetically inactivated JONs; rather, we observed a decrease in wingbeat frequency relative to control flies, and only at 100 cm/s and lower wind speeds (Figure 4E, yellow; all p<.05, Holm-corrected). In contrast, we observed a similar decreasing trend in relative wingbeat amplitude as wind speeds increased across all three experimental groups here (Figure 4F). Although we found a steeper decrease in wingbeat amplitude as wind speed increased in flies with glued antennal joints relative to flies with intact JONs (Figure 4F, blue; p=.012, linear-mixed model), we observed no differences between flies with intact JONs and flies with either silencing manipulation at any wind speed (all p>.05, Holm-corrected).

### Flies actively position their antennae in response to varying optic flow speed during flight

To understand how progressive visual information affects antennal positioning in the absence of frontal airflow, we presented flies with a vertical bar pattern moving at rates of 0-50 cm/s with no airflow (Figure 5A, STAR Methods). In flying flies, we observed an active outward movement of the antennae at a high (35 cm/s) optic flow speed compared to a low optic flow speed (5 cm/s), but in nonflying flies, antennal angle was approximately unchanged over this speed range (Figure 5B). When comparing no optic flow (0 cm/s) to the lowest optic flow speed we tested (5 cm/s), we saw a significant decrease in active inter-antennal angle in flying flies but not nonflying flies (Figure 5C; p<.05 for flying flies, paired t-test). To measure optic-flow-speed-dependent changes in antennal responses, we quantified movements of the antennae relative to the lowest optic flow speed (5 cm/s). Nonflying flies did not significantly change their antennal position at any optic speed except 35 cm/s (Figure 5D, left; p<.01 at 35 cm/s, p>.05 for rest, Holm-corrected one-sample t-tests). However, in flying flies, we observed clear outward active movements of the antennae with increasing optic flow (Figure 5D, right; p<.001, linear-mixed model), and baseline-relative antennal position was significantly different from zero at all optic flow speeds (all p<.05, Holm-corrected one-sample t-tests).

**Figure 5.**
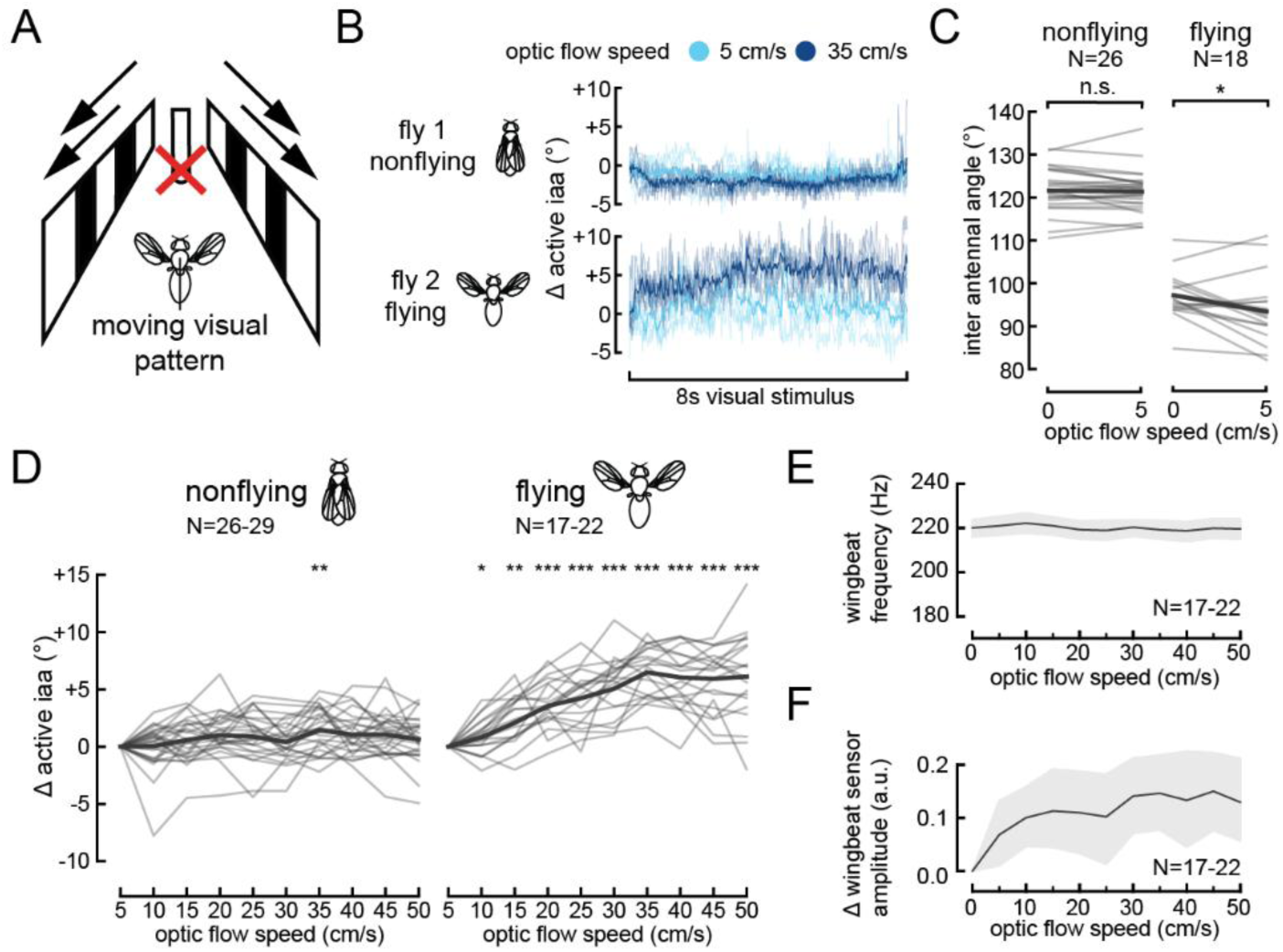
Flying flies increase inter-antennal angle with increasing optic flow rate. A) Schematic of the optic flow experiment. The airflow tube was present with no airflow delivered. B) Average active (second segment) inter-antennal angle in response to progressive optic flow moving at 5 cm/s (light blue) and 35 cm/s (dark blue) for an example nonflying (top) and flying (bottom) fly for a single 8 sec trial. Shaded traces represent the average across 5 trials; transparent traces represent individual trials. C) Absolute active inter-antennal angle at 0 cm/s and 5 cm/s optic flow in nonflying flies (left, p=.27) and flying flies (right, p=.025). Average across all flies is represented by shaded traces, while individual flies are represented by faded traces. D) Average individual (grey) and cross-fly (black) active inter-antennal angle across all optic flow speeds (excluding zero), baseline-subtracted relative to the lowest speed (5 cm/s) for nonflying (left) and flying (right) flies. The active antennal response of flying flies is significantly larger than nonflying flies (β=.123, p<.001) and greater than zero at all non-baseline optic flow speeds (all p<.05). E) Wingbeat frequency (top) and relative wingbeat amplitude (bottom) across all optic flow speeds.

Across optic flow speeds, we did not observe substantial changes in wingbeat frequency (Figure 5E). However, we found that wingbeat amplitude grew slightly with increasing optic flow (Figure 5F). This contrasts with our observations in flies presented with only varying airflow, where we found that wingbeat amplitude consistently decreased as airflow increased (Figure 2G and 4F). These results suggest that steady-state differences in airflow and progressive optic flow have inverted effects on wingbeat amplitude in open-loop (tethered flight), with faster optic flow driving increases in wingbeat amplitude and faster airflow decreasing wingbeat amplitude.

### Frequency-dependent responses to dynamically changing airflow and optic flow

To understand how mechanosensory and visual information are combined to influence active sensation by the antennae, we presented both wind and progressive optic flow stimuli synchronously (Figure S2, STAR Methods). We delivered wind and optic flow as sinusoidal stimuli at three frequencies (0.3, 1.3, 2.3 Hz), oscillating around a set baseline value simulating forward flight. In natural settings, optic flow speed depends strongly on the geometry of the visual environment (e.g., distant objects traverse the retina more slowly than objects closer to the observer), and thus there is no natural correspondence between optic-flow speed and wind speed. Thus, we measured steady state antennal responses to wind speed (Figure S4A) and optic flow (Figure S4B) to determine oscillatory ranges (100-200 cm/s for wind, 5-35 cm/s for visual) and baseline values (150 cm/s wind, 20 cm/s visual) for both oscillating stimuli. We presented one stimulus oscillating from its lowest to highest range in a sinusoidal pattern while holding the other modality at its baseline value (and in the opposite manner with the second stimulus). Additionally, we presented both wind and visual stimuli oscillating simultaneously within their respective ranges.

To confirm we were not saturating the antennal motor system with our stimulus (i.e. active antennal movements reached their maximum range), we first presented each stimulus type at half amplitude (125-175 cm/s for wind, 12.5-27.5 cm/s for visual) and full amplitude (100-200 cm/s for wind, 5-35 cm/s for visual) (Figure S4C). We observed that wind oscillation-dependent movements scaled approximately linearly across oscillation frequencies in nonflying flies (Figure S4D, blue), while visual oscillation and wind plus visual oscillation-dependent movements exhibit nonlinear properties (however we note that visual responses were, in general, low in nonflying flies; Figure S4D, red and purple). For flying flies, all three stimulus combinations elicited mildly nonlinear responses (and highly variable responses for vision-only), but generally scaled (roughly) by a factor of 2, indicating that the stimuli were not saturating the motor response (Figure S4E).

In response to oscillating airflow and optic flow, we observed changes in active antennal responses in both nonflying and flying flies that depended on the frequency of oscillation and the sensory modality that was oscillating (Figure 6A). Antennal motions in nonflying flies were smaller across all oscillation frequencies and oscillatory conditions than in flying flies (compare gains in Figure 6B vs. 6D). This was especially true when optic flow speed alone was oscillating, with nonflying flies exhibiting near-zero active movements (Figure 6A and 6B, red). Notably, these movements in response to optic flow oscillations also lagged in phase when compared to the other stimulus conditions (Figure 6A and 6C, red).

**Figure 6.**
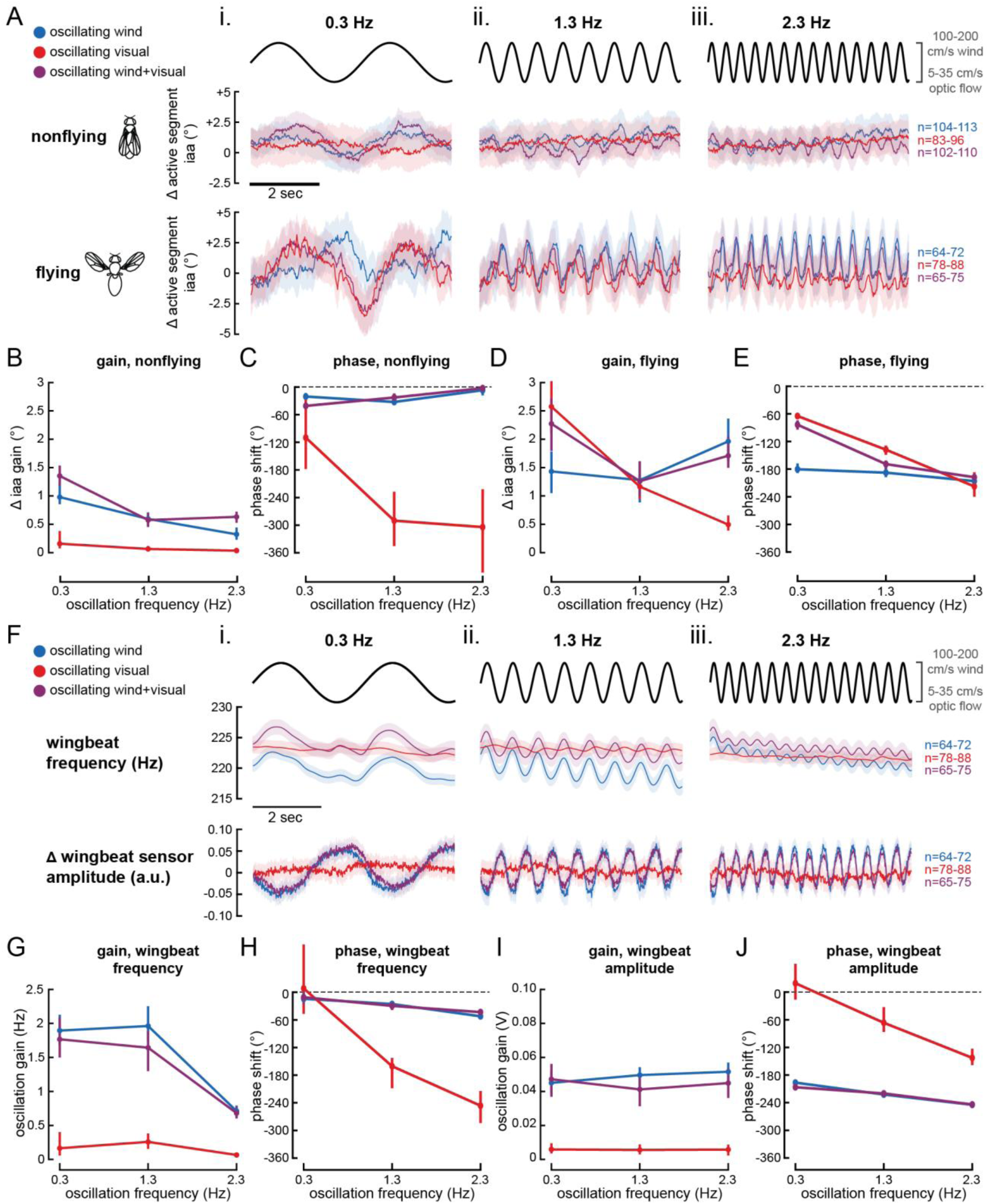
Active antennal responses to oscillating combinations of airflow and optic flow reveal flight-gated nonlinear integration. A) Average baseline-subtracted active segment inter-antennal angle over a 6 sec period in response to oscillating wind speed (with constant 20 cm/s optic flow, blue, N=14-15 flies), oscillating optic flow (with constant 150 cm/s airflow, red, N=15 flies), or synchronous oscillation of wind speed and optic flow (purple, N=15 flies) for nonflying (upper middle) and flying (bottom) flies. Black lines (top) represent time-dependent location of stimulus oscillation on an arbitrary y-scale. Shaded regions represent one standard error of the mean. B) Gain of active antennal movements across oscillatory conditions in nonflying flies. Error bars are generated by a 100-epoch Monte Carlo estimation with replacement at each frequency. C) Phase of active antennal movements relative to time-dependent phase of stimuli presentation (black, top of (A)). Error bars are generated by same method as (B). D) Same as (B), in flying flies. E) Same as (C), in flying flies. F) Wingbeat frequency response (upper middle) and baseline-subtracted wingbeat amplitude (bottom) relative to oscillating stimuli (black, same as (A)) in flying flies. Dataset originates from flying flies in (A). G) Gain of wingbeat frequency response across oscillatory conditions. Error bars are generated by a 100-epoch Monte Carlo estimation with replacement at each frequency. H) Phase of wingbeat frequency response relative to time-dependent phase of stimuli presentation (black, top of (F)). I) Gain of baseline-subtracted wingbeat amplitude across stimulus oscillation frequencies. J) Phase of baseline-subtracted wingbeat amplitude across stimulus oscillation frequencies.

Flying flies performed larger antennal movements than nonflying flies, and these movements depended strongly on the oscillation frequency (Figure 6A, bottom). At 0.3 Hz, the gain (Figure 6D) and phase (Figure 6E) of visually driven movements (Figure 6Ai, bottom, red) were nearly identical to that of the responses to the combined (wind and visual) cue (Figure 6Ai, bottom, purple). In contrast, at 2.3 Hz, the visually driven movements were much smaller (Figure 6D, red) and wind-oscillation-dependent movements (Figure 6Aiii, bottom, blue) were nearly identical to the responses to the combined cue (Figure 6Aiii, bottom, purple). As oscillation frequency increased, the phase lag (Figure 6E) of wind-driven dependent movements remained steadily antiphase (blue), suggesting a flat mechanosensory gain of -1, while visually driven (red) and wind-plus visually driven (purple) movements lagged slightly in phase at the lowest frequency, and trended towards antiphase. Critically, these results indicate that for tethered, flying flies, low-frequency movements were dominated by vision, whereas high-frequency antennal movements by mechanosensation, crossing over at the central frequency we tested. In contrast, for nonflying flies, antennal responses followed mechanosensory cues across all frequencies of sensory oscillations.

We also observed differences in both wingbeat frequency and wingbeat amplitude depending on the type of oscillating stimuli and oscillation frequency (Figure 6F). The gain (Figure 6G) and phase (Figure 6H) of wingbeat frequency oscillations closely matched each other in the wind oscillation (blue) and wind-plus visual-oscillation (purple) conditions, with a significant decrease at 2.3 Hz. However, the wind-plus visual-oscillation condition (purple) tonically increased wingbeat frequency at all oscillation frequencies compared to wind-only (blue) oscillations (Figure 6F, ii); we note that this tonic increase does not manifest as a change in the frequency response gain or phase (Figure 6G). With oscillating visual stimuli, the magnitude of wingbeat frequency changes is near zero, and phase lag increases dramatically as oscillation frequency increases (Figure 6G-H, red).

Similarly, we measured minimal changes in wingbeat amplitude in response to oscillating visual stimuli (Figure 6I, red) with phase lag increasing significantly as oscillation frequency increases (Figure 6J, red). In the wind-only (blue) and the wind-plus visual-(purple) oscillation conditions, we observed similar oscillations in wingbeat amplitude responses (Figure 6I). The phase lag of these two conditions mirrors that of wingbeat frequency, albeit in antiphase.

## DISCUSSION

### Visual environment modulates active antennal sensing

In our study, we directly investigated how airflow and wide-field visual stimuli control the movements of (and by extension, sensation by) the multimodal antennae of *Drosophila melanogaster.* We first observed more frontal positioning of the antennae as frontal airflow increased in darkness (Figure 2D). The fly’s response inverted in the presence of a static visual grating, with antennae position shifting backwards with increasing frontal airflow (Figure 2E). These inversions in behavioral responses to broad changes in visual environment are similar to previously reported work. For instance, in one closed-loop tethered experiment^74^, flies oriented downwind in response to airflow in an enclosed, dark environment. In a separate closed-loop tethered study, flies instead exhibited an inverted response, orienting upwind in the presence of airflow with some ambient light present^49^. Light levels can have a profound effect on behavior in other insect species like the honeybee *Apis mellifera*, which halt flight in dark conditions^75^. Our results support the conclusion that both the presence and content of the visual environment strongly influence flight behavior. Notably, the presence of progressive visual motion resulted in an opposite wingbeat amplitude response compared to a static visual environment.

In response to visual motion, we consistently observed an increase in the magnitude of antennal movements in flying flies compared to nonflying flies. This was true with steady-state optic flow in the absence of airflow, where only flying flies performed significant visually driven antennal movements (Figure 5). Additionally, we observed an increased gain across all oscillatory sensory conditions in flying flies as compared to their nonflying counterparts (Figure 6A-E). However, we did not observe flight-dependent changes in antennal responses to airflow in different visual environments, in the absence of visual motion (Figure 3C). This suggests that increased magnitude of antennal movements during flight requires the presence of visual motion. One potential explanation for this phenomenon is the direct modulation of upstream visual motion circuitry during flight by octopaminergic neurons^76–78^, which might increase the salience of visual information integrated by antennal motor circuits. It is also possible that a nonlinear interaction between visual motion information and an internal representation of flight in premotor antennal circuits exists; evidence that visual input modulates antennal control circuitry in other flying insects, such as the hawkmoth (*D. nerii*)^79^ and honeybee (*A. mellifera*)^80^, may support this explanation. In this scenario, reafferent feedback from wingbeats detected by Johnston’s Organ Neurons (JONs)^47,67,81^ could be integrated directly by premotor circuitry. Additionally, flight state information from ascending neurons may directly convey behavioral states to central brain premotor regions, similar to those found in grooming and walking behaviors^82^. Further investigation through connectomics analyses and *in vivo* physiology in behaving animals may enable us to discern how visual motion information is relayed to antennal motor circuits in a state-dependent manner.

### Control of antennal position at the onset of flight

In our study, we observed a steady-state frontal positioning of the antennae during flight, consistent with previous work^26,65,68–70^ (Figure 3A-B, 4B-D). This flight-state positioning is invariant between flies in darkness and those presented with a static grating (Figure 2E, left) in the absence of airflow, which is surprising based on our observation that they perform different antennal movements as airflow increases (Figure 2E, right). We also found that silencing JON input both mechanically and optogenetically shifts this flight-state positioning to a more extreme frontal position across all wind speeds (Figure 4B, 4D). This result is in contrast to previous work in hawkmoths, where mechanical silencing produced no difference in flight-based forward antennal positioning^65^. Rather, in this previous study, the Böhm’s bristles (hairs positioned between cuticular segments thought to aid in proprioception) controlled flight-based positioning of the antennae. Additionally, silencing hawkmoth and honeybee JON activity with glue halts wind speed-dependent antennal movements^80,83^. Surprisingly, our results do not support the conclusion that airflow-dependent changes to antennal positioning in flight is mediated primarily by JONs (Figure 4D). However, our finding that wind speed-dependent responses persist in flies with silenced JONs (mechanically and optogenetically) suggests that a sensor analogous to Böhm’s bristles, such has hair plates detecting relative position between the first and second segments (similar to those recently described in *Drosophila* legs^84^), may contribute to antennal positioning in *Drosophila*. Nonetheless, we observed a steady dysregulation of inward antennal positioning in flies with silenced JONs, largely independent of wind speed (Figure 4B), suggesting that a feedforward motor program may be involved in this behavior.

Unlike with antennal movements, whose airflow-dependent responses appear to be independent of JON sensation, we observed a clear effect of blocking antennal sensation on wind speed-dependent wingbeat frequency (Figure 4E). Markedly, wingbeat frequency is mostly unaffected by optogenetically silencing the JONs, but mechanically blocking the passive joint with glue resulted in significant increases in wingbeat frequency as wind speed increases. To optogenetically manipulate JONs, we used a genetic driver line (*nan*-GAL4) that labels all chordotonal organs^72,73^ (stretch-receptive sensors also located in the femur, haltere, and wings). Silencing these additional sensory cells outside the antenna could have contributed to a decrease in wingbeat frequency compared to the response in flies with glued antennae. However, the two antennal sensory manipulations (mechanical and optogenetic silencing) produce similar changes in antennal response at flight onset and in response to varying airflow, supporting the conclusion that antennal sensation is crucial for guiding these active behaviors.

### Mechanisms of visual and mechanosensory integration

One caveat of our findings is that the open-loop tethered preparation does not allow the animal’s intended changes in speed to change the resulting wind flow patterns on the antenna, nor the perceived optic flow. Thus the flight control system is operating in open loop (Figure 1D), and because the visuomotor transform is thought to act like an integrator for forward flight speed control^51^, the small effect of visual oscillations on wing kinematics may arise, at least in part, due to this lack of reafferent feedback, which is known to heavily attenuate sensorimotor responses^85^.

When we provided nonflying flies with oscillatory airflow and optic flow, antennal movements generally mirrored the responses to mechanosensory oscillations alone across all frequencies (Figure 6B and 6C). This broad-band mechanosensory winner-takes-all computation is illustrated in Figure 7A. In contrast, we observed an unexpected nonlinear effect of oscillation frequency on antennal movements in flying flies; as the oscillation frequency increases, antennal movements in response to synchronous airflow and optic flow oscillations (Figure 6D,E, purple) shift from mirroring the antennal response to visual oscillations (Figure 6D,E, red) to mirroring the antennal response to oscillating wind stimuli (Figure 6D,E, blue). This frequency-dependent “winner-takes-all” effect between visual information at lower frequencies and mechanosensory information at higher frequencies (Figure 7B) is similar, for instance, to hawkmoth flight tracking of food source (*M. stellatarum*)^86^ and head stabilization during body rolls (*D. nerii*)^87^. Additionally, our findings may be analogous to multimodal integration of visual and haltere signals during turning behavior in fruit flies, where slower turns rely on visual information and faster turns use mechanosensory feedback from the angular velocity encoding halteres^88^. Mechanical transduction is lower latency than visual transduction making it more suitable for rapid (high frequency) responses. Although the visuomotor response may dominate at low frequencies^51,89^, it is also slower than mechanosensory feedback due to phototransduction^90^. This makes visual information inherently less useful as motion frequency increases, evidenced by the increasing phase lag we observed (high phase lag feedback can, if not attenuated, destabilize closed-loop dynamics). Thus, the faster mechanosensory pathway may guide appropriate antennal positioning at higher frequencies, as is likely essential for highly variable environments. Altogether, our results demonstrate previously uncharacterized multisensory control principles for active antennal sensation and pave the way towards understanding the precise neural circuits responsible for performing these computations.

**Figure 7.**
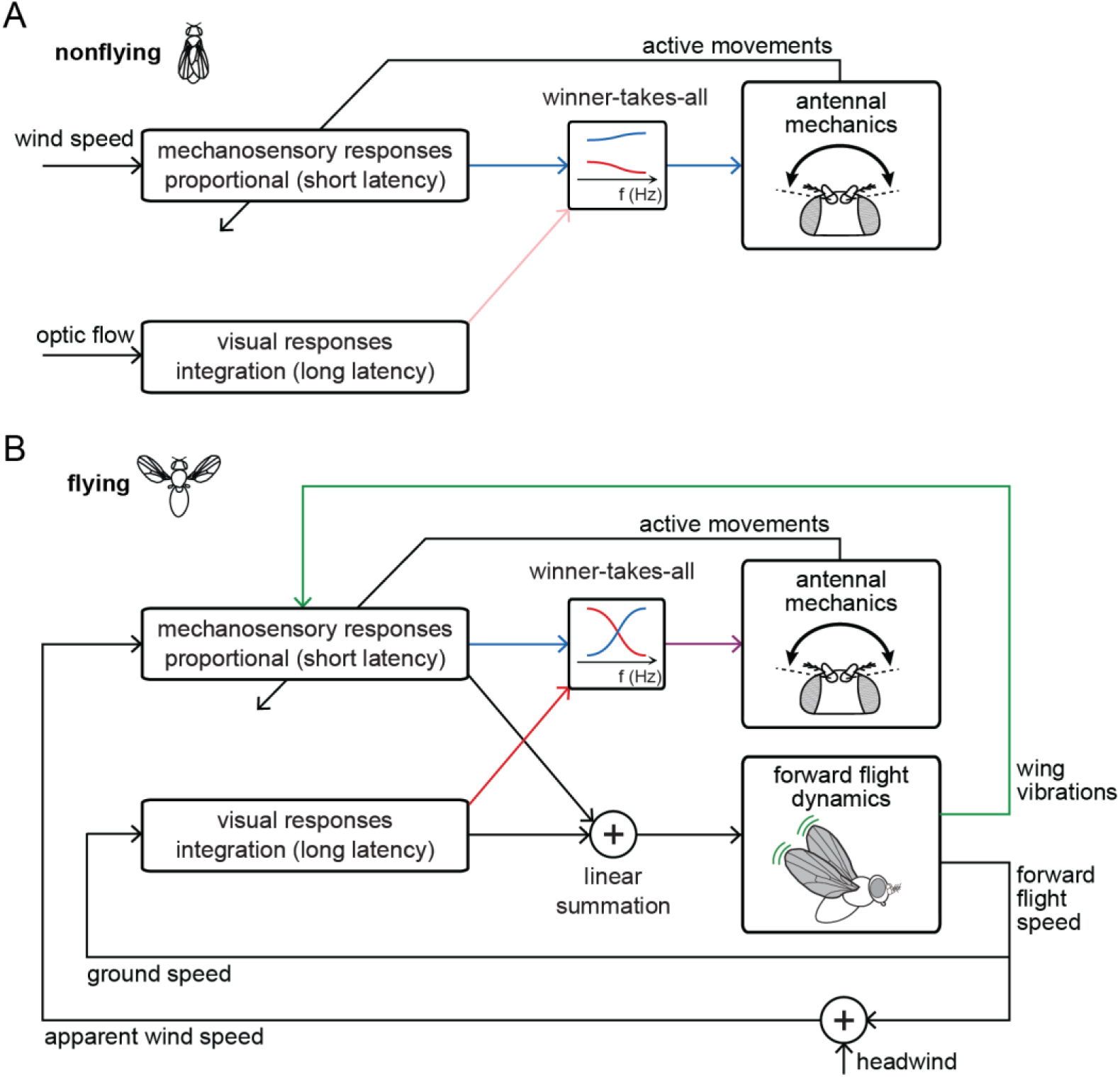
Integration of mechanosensory and visual cues for antennal control is nonlinear and state-dependent. A) For nonflying flies, mechanosensory cues created by oscillating wind speed elicited robust antennal responses. Time-varying optic flow stimuli, by contrast, elicited weak antennal responses that were overridden by responses to mechanosensory cues when both were present. B) A simplified control theoretic model illustrates the potential role of active antennal movements during forward flight. Based on prior work^51^, forward flight speed control is thought to be well approximated by the linear summation of a proportional, lower-latency mechanosensory response and an integral, longer-latency visual response (lower multisensory feedback loop in the diagram). Our work shows that antennal movements were regulated in a frequency-dependent, “winner-takes-all” manner, where low-frequency antennal motions are dominated by visual feedback and high-frequency antennal motions are dominated by mechanosensory feedback (upper multisensory feedback loop). We hypothesize that this active sensing control, illustrated by the diagonal arrow through the mechanosensory response block, is modulatory in nature, altering antennal neuromechanical processing of wind speed. This suggests that the dynamics of forward flight control emerge from a neuromechanical system that includes inner-loop sensor modulation.

## ACKNOWLEDGEMENTS

We would like to thank Peter Polidoro (Janelia/HHMI Research Campus) for designing the wingbeat sensor and John Fellenstein (Technical Supervisor III, Division of Science Machine Shop, Vanderbilt University) for machining assistance. Critical support and facilities for photodiode instrumentation development and testing were provided by Anupam Kumar and the Wond’ry Making and Design spaces at Vanderbilt University. We also thank Bradley Dickerson, Jessica Fox, and Amy Streets for helpful feedback. This research was supported by the NIH through a BRAIN Initiative R00 NS114179 (M.P.S.) and BRAIN Initiative U01 NS131438 (N.J.C. and M.P.S.). Additionally, stocks obtained from the Bloomington Drosophila Stock Center (NIH P40OD018537) were used in this study.

## AUTHOR CONTRIBUTIONS

Conceptualization, K.M.M., N.J.C., and M.P.S.; Methodology, K.M.M., N.J.C., and M.P.S.; Formal Analysis, K.M.M.; Investigation, K.M.M.; Writing-Original Draft, K.M.M., N.J.C., and M.P.S.; Writing-Review and Editing, K.M.M., N.J.C., and M.P.S.; Visualization, K.M.M., N.J.C., and M.P.S.; Supervision, M.P.S.; Funding Acquisition, N.J.C and M.P.S.

## STAR METHODS

### KEY RESOURCE TABLE

| REAGENT or RESOURCE | SOURCE | IDENTIFIER |
| --- | --- | --- |
| Chemicals, peptides, and recombinant proteins |  |  |
| all-trans retinal | Sigma-Aldrich | Cat #R2500 |
| Deposited data |  |  |
| Raw and analyzed data | This paper | Available upon publication through an open data publishing platform |
| Experimental models: Organisms/strains |  |  |
| *D. melanogaster*: Canton-S | Dickinson Lab | N/A |
| *D. melanogaster*: w[*]; P{y[+t7.7] w[+mC]=UAS-GtACR1.d.EYFP}attP2 (UAS-GtACR1) | Bloomington Drosophila Stock Center | RRID: BDSC_92983 |
| *D. melanogaster*: w[*]; P{w[+mC]=nan-GAL4.K}2 (nan-GAL4) | Bloomington Drosophila Stock Center | RRID: BDSC_24903 |
| Software and algorithms |  |  |
| MATLAB version 2023b | MathWorks | RRID: SCR_001622 |
| Psychophysics Toolbox | https://psychtoolbox.org/ | RRID: SCR_002881 |
| Python version 3.10.11 | Python Software Foundation | https://www.python.org/downloads/release/python-31011/ |
| seaborn | https://seaborn.pydata.org/ | RRID: SCR_018132 |
| SciPy | http://www.scipy.org/ | RRID: SCR_008058 |
| statsmodels | http://www.statsmodels.org/ | RRID: SCR_016074 |
| emd | https://emd.readthedocs.io/en/stable/ | N/A |
| Custom code for project | This paper | https://github.com/suverlab/MillsEtAl2025 |

## RESOURCE AVAILABILITY

### Lead contact

Any requests for resources or further information should be directed to and will be fulfilled by the lead contact, Marie P. Suver.

### Materials availability

No new materials or reagents were generated in this study.

### Data and code availability

We would like to thank the developers of seaborn^91^, SciPy^92^, statsmodels^93^, and emd^94^ for their contributions to free and open-source software. Data and code generated in this study will be made available via Dryad and Github at time of publication.

## EXPERIMENTAL MODEL AND SUBJECT DETAILS

All experiments used adult female *Drosophila melanogaster* between 3 to 7 days post-eclosion. All flies were raised on standard yeast-cornmeal medium at 25° C on a 12-hour light-dark cycle. Experimental genotypes for each figure are listed in the table below. Parental genotypes and RRIDs are available in the Key Resource Table.

### Fly genotypes used in experiments

|  |  |
| --- | --- |
| Figure 2 | *Canton-S* |
| Figure 3 | *Canton-S* |
| Figure 4 | *Canton-S>UAS-GtACR1; nan-GAL4>UAS-GtARC1* |
| Figure 5 | *Canton-S* |
| Figure 6 | *Canton-S* |
| Figure S1 | *Canton-S* |
| Figure S4 | *Canton-S* |

## METHOD DETAILS

### Behavioral apparatus

To present airflow, humidified air was sent through a tube (McMaster 89895K657) positioned approximately 5 cm anterior to the fly. Wind speed was controlled by a mass flow controller (Dakota Instruments 6AGC1AL55-08AB) and toggled on and off by a solenoid valve (Lee Company LHDA1233115H) controlled by a custom MATLAB script. To measure and calibrate wind speed, we used a calibrated hot wire anemometer (Dantec MiniCTA with 55P11 probe). Additionally, we used a manual wind puffer to elicit flight, with an output tube positioned dorsally to the primary airflow tube. Puff occurrences were measured by an airflow sensor (Honeywell Sensing Solutions AWM3300V).

To block external light, we created a 24”x24” lightproof box with foamcore walls and ceiling mounted on a 1/4”-20 breadboard (Thorlabs MB2424). To present unified optic flow, two 308.448 x 93.43 mm (2880 x 864 pixels) 120 Hz LCD screens were places parallel to the fly on both left and right side at a distance of 87 mm. Visual stimuli presented were bilaterally and vertically symmetric, so for practical purposes the right screen displayed a vertically flipped view of the left screen. All vertical bars within the visual gratings were 35.343 mm across (330 pixels), totaling approximately 22 degrees within the fly’s field of vision when an edge is directly adjacent to the fly. Reflections of light on the LCD screens were minimized with custom-fitting anti-glare screen covers.

We created all visual stimuli in PsychoPy, and all visual stimuli were presented using Psychophysics Toolbox in a secondary MATLAB instance connected to the primary MATLAB instance through TCP/IP protocol. To synchronize visual stimuli and wind stimuli presentation, we built a custom photodiode-based circuit that would trigger the primary instance of MATLAB to both begin data acquisition and initiate the wind stimulus upon detection of a localized, time-synchronized black-white oscillation on the LCD screen out of view of the fly (Figure S2). The photodiode sensor was placed facing a corner of the visual display outside the field of view of the animal; the sensor produced a voltage proportional to the intensity of light detected. When the voltage output from the sensor increased above a threshold of 2.5 V, it results in a trigger signal sent to the computer/data acquisition (DAQ) object, which would instantaneously trigger airflow.

All voltage data was collected and transmitted at a sampling rate of 20 kHz through a 4 AO PCIe card (National Instruments PCIe-6323) attached to two BNC breakout boards (National Instruments BNC-2110).

### Experimental design

In the experiments shown in Figure 2-4 and Figure S1, each fly experienced 10 airflow trials either increasing from 0 cm/s to 300 cm/s (5 trials) or decreasing from 300 cm/s to 0 cm/s (5 trials). Airflow was presented in 50 cm/s steps over 42 sec, with 6 sec of steady state airflow for each wind speed. The order of increasing and decreasing trials was pseudo-randomized for each flight. For each wind speed, averages were taken over the last 4 sec as the mass flow controller did not adjust instantaneously. 6 sec of pre-trial at the initial wind speed (with the solenoid valve off, to prevent airflow) was fed to the mass flow controller to allow for adjustment and settlement at the correct speed before the trial began. Additionally, at 0 cm/s wind speed the solenoid valve was shut off to prevent airflow.

In the experiments shown in Figure 5, each fly experienced 60 optic flow trials ranging from 0-50 cm/s in 5 cm/s increments (5 trials at each optic flow speed, pseudo-randomized order). Each trial consisted of an 8 sec period of constant optic flow. For each optic flow speed, averages were taken over the last 6 sec of each trial. The interlude between trial periods consisted of a non-moving static grating, equivalent to the 0 cm/s optic flow condition. The solenoid valve for airflow was shut off for the entirety of these visual stimulus only experiments.

In the experiments shown in Figure 6, each fly experienced 36 trials at 3 oscillatory frequencies (0.3, 1.3, and 2.3 Hz) with wind speeds ranging from 100-200 cm/s and optic flow speeds from 5-35 cm/s. Trials were presented in 12 blocks, with each frequency represented once pseudo-randomly organized within each block. Wind and visual stimuli were presented synchronously during the experiment. If oscillations were not applied to a stimulus type (wind or visual) in a condition, the non-oscillating stimulus was held at a constant value in the middle of its oscillatory range (150 cm/s for wind, 20 cm/s for visual). Trials were 16 sec in length, with 2 sec before each trial used for synchronization of visual and wind stimuli through the photodiode circuit and 1 sec after each trial used to shut off airflow (via solenoid valve) and the camera recording. Pre-trial and post-trial data was omitted during post-processing. Each trial consisted of 4 sec of baseline presentation, which consisted of both wind and visual stimuli being held at the middle of their respective oscillatory ranges (150 cm/s for wind, 20 cm/s for visual). The following 12 sec consisted of stimuli oscillation or stimuli being held in the middle of its oscillatory range depending on the condition.

To confirm our accurate manipulation of wind speed at high frequencies, we performed directed calibration experiments using the hot wire anemometer (Dantec MiniCTA with probe). The anemometer was placed in the position of the tethered fly during experiments. First, we performed a sweep of all wind speeds in our experimental range (0-300 cm/s in 1 cm/s increments) and collected the average anemometer value at each speed (Figure S3A). Using this, we fit a high-order polynomial to the data (overfitting was not a concern, as we did not extrapolate nor leave the original experimental range) and found the inverse of the polynomial to use as a transfer function for anemometer values to wind speed. When applying the same voltage oscillation (± 1.1 V) from a baseline (∼2.49 V, equivalent to 150 cm/s) at 0.1 Hz and 1.1 Hz, a smaller wind speed oscillation amplitude was observed at the higher frequency (Figure S3B). This was observed across frequencies between 0.1 and 1.1 Hz and multiple oscillation voltages (± 0.55 V, ± 1.1 V, and ± 2.2 V), with negative linear relationships between oscillation frequency and oscillation amplitude consistently found (Figure S3C). With the knowledge that oscillation voltage does not equivalently scale oscillation amplitude, we hand-tuned (by adjusting oscillation voltage) oscillation amplitude across various frequencies to be equivalent to ± 50 cm/s (Figure S3D).

We also measured the delay between the command sent to mass flow controller and the arrival of the wind stimulus. To account for this in our stimulus timing, we experimentally measured the phase of the wind stimulus at each frequency. This phase delay was then applied to the visual stimulus so that both wind and visual stimuli would be synchronized when oscillating together.

### Fly preparation

Flies were first anesthetized on ice and placed on a metal plate liquid-cooled by ice-cold water flowing through a peristaltic pump. We then dissected all 6 legs at the femoral-trochanter joint to ensure no grooming of antennae or wings would occur over the course of the experiment. Ultraviolet-curing glue (TOPCHASE B08YMTFM7D) was then placed in the gap between head and thorax to fix the head in place during experiments. The rigid tether was then lowered at an 85-degree angle relative to the rostral-caudal axis of the fly and fixed to the mid-thorax using ultraviolet-curing glue. The tethered fly was then placed into the behavioral apparatus at a 60-degree angle relative to ground.

All flies were rigidly tethered and head-fixed using the same protocol, except for dead flies (Figure 2A-B) and those with glued active-passive antennal joints (Figure 4B-E). Dead flies were frozen at approximately -25° C for 5 minutes before tethering, following a method similar to previous work^49^. We confirmed that the flies did not reawaken after this freezing period. To glue the active-passive antennal joint, we applied a small drop of ultraviolet-curing glue to the medial edge of the second (pedicel) and third (funiculus) segments of each antenna. To confirm successful application, we nudged each arista with a paintbrush after the glue was cured to verify the second segment moved in tandem with the third segment.

### Antennal tracking

To record antennal movements, the fly was illuminated from above by infrared light (850 nm) emanating from a 10x2 grid of infrared LEDs (ORDRO LN-3 Studio IR Light Accessory). Antennal movements were measured using a camera (Allied Vision Guppy Pro F-031 with InfiniStix 44mm/3.00x lens) rotated ten degrees backwards from perpendicular to ground. Videos were recorded at 60 frames per sec at a resolution of 640x480 pixels.

We used DeepLabCut^61^ (version 2.3.6) to track antennal movements over time. The network was trained using 670 frames from 67 flies across genotypes and experimental designs. We tracked 20 points on the head and antennae of the fly, including 6 static points on the head to determine a head axis, 3 on each third antennal segment (including the arista) and 4 on each second antennal segment. We used a ResNet-50 neural network, trained over 500,000 iterations with a training-to-test fraction of 0.8. The trained network demonstrated a training error of 1.94 pixels and a testing error of 2.24 pixels, respectively.

### Wingbeat data collection

To gather information on wingbeat frequency and amplitude, infrared light reflected off the beating wings was collected via a liquid light guide (Thorlabs LLG3-4Z) and fed into a custom circuit board. The signal was smoothed via a second-order bandpass Butterworth filter (cutoff frequencies: 150-250 Hz) followed by a second-order Savitsky-Golay filter (window size: 47 samples). We determined if the fly was flying or quiescent by taking the absolute value of a Hilbert transformation of the processed wingbeat signal followed by a second-order lowpass Butterworth filter (cutoff frequency: 6 Hz); this was then compared to a threshold value. Instantaneous frequency was estimated from the processed wingbeat signal using an amplitude-normalized Hilbert transformation^95^.

To assess whether our photodiode-based measurement of wingbeat amplitude correlated with true amplitude measured using video recording, we tracked the wing envelope using a camera (Allied Vision Guppy Pro F-031 with InfiniStix 94mm/0.45x) mounted above the fly (Figure S1A). For our photodiode-based measurement of instantaneous wingbeat amplitude, we took the maximum amplitude values of the original wingbeat signal over 100 sample (0.05 sec) window periods (Figure S1B). For our camera-based measurement, we tracked the tip and base of the leading edge of the fly’s wings, as well as the midline of the fly using DeepLabCut^61^ (Figure S1C). We trained a ResNet-50 neural network over 500,000 iterations on 330 frames, sourced from 22 videos from 11 flies (2 per fly). The training-to-test fraction for this network was 0.8. The trained network demonstrated a training error of 2.03 pixels and a testing error of 3.92 pixels, respectively. To compare the camera measurement with our photodiode measurement, we used the combined left- and right-wing angle from leading edge to fly midline (‘wingstroke angle’). We observed a negative linear relationship between wingstroke angle and wingbeat sensor amplitude (Figure S1D, r=-.7754, r^2^=.6013, p<.001) and accordingly sign-inverted our wingbeat sensor amplitude so that lower photodiode-based, raw measurements display as larger wingbeat amplitudes.

### Optogenetic inactivation

All flies used in optogenetic experiments were placed on food saturated with all-trans retinal 24-48 hours prior to tethering. To create the retinal saturated food, 50 µL all-trans retinal (Sigma-Aldrich R2500) was mixed with 0.75 g rehydrated potato flakes and placed on top of standard fly-food mixture. For GtACR1^71^ inactivation experiments, we used 505 nm cyan light (Thorlabs, Driver: LEDD1B, LED: M505F3) calibrated at an intensity of 7.00 mW/cm^2^. Light was delivered by a raw fiber optic patch cable (Thorlabs M118L02) positioned dorsally to the fly, with intensity measured at the location of the tethered fly.

## QUANTIFICATION AND STATISTICAL ANALYSIS

All data was collected in MATLAB 2023b and analyzed using custom Python scripts. Alpha values were set at .05 for all statistical tests, with * representing p<.05, ** representing p<.01, and *** representing p<.001.

To analyze active and passive antennal movements, we computed the angles for both the second and third antennal segments relative to a head axis midline using the points in Figure 1C. Inter-antennal angle for active and passive antennal movements was calculated by adding the angle of the left and right antenna for each segment respectively.

Across all behavioral groups, the Kolmogorov-Smirnov test (scipy.stats.kstest) was used to confirm normality of data.

For most statistical analyses (Figure 2D-G, 3C, 4B-F, 5D), we used a linear mixed-effect model (statsmodels.formula.api.mixedlm) with post-hoc tests to determine individual differences between groups at each stimulus level. All but one model included the stimulus (wind speed or optic flow speed) as a continuous variable (in increments of 50 cm/s and 5 cm/s respectively) and the conditional groups (e.g., darkness v. static grating, flying v. nonflying, and JON silencing) as categorical variables. Both stimulus speed and conditional group were considered fixed effects (estimated using restricted maximum likelihood, REML); individual flies were included as random intercepts. The model was specified as follows:

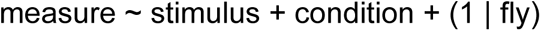

Where measure is inter-antennal angle, wingbeat frequency, or wingbeat amplitude.

For Figure 4B, we used two categorical variables: flight state and condition. Given the relative simplicity of the model, we included random slopes per fly for flight state to increase model accuracy. The results of all linear-mixed models and their descriptive equations can be found in Table S1-4.

To test for linearity and homoscedasticity, we generated residuals vs. fits plots and looked for abnormality in variance. While some “pinching” was observed near zero in models using baseline-subtracted data (which was expected, given reduced variability near baseline), we did not observe any major abnormalities. We also checked for normality using Q-Q plots, and did not detect any substantial violations of assumptions.

Post-hoc analysis was performed through model-derived estimated marginal mean (EMM) contrast comparisons across conditional groups at each stimulus speed (wind speed or optic speed). To control Type I error, we then performed a Holm-correction (statsmodels.stats.multitest.multipletests) on generated p-values on a per model basis.

Other analyses performed included independent Student’s t-tests without correction (Figure 2D-E), paired Student’s t-tests (scipy.stats.ttest_rel, Figure 3C, 5C), one-sample Student’s t-tests with Holm-correction (scipy.stats.ttest_1samp, Figure 5D), and a linear regression (scipy.stats.linregress, Figure S1D).

In Figure 6, we performed a Fast Fourier Transform (scipy.fft.fft and scipy.fft.fftfreq) on all average traces to obtain oscillation gain and phase lag information (Figure 6B-E, 6G-J, S4D-E). We normalized the fft output by dividing it by a normalized-to-one unitless sinewave in phase with the oscillating stimulus. We calculated oscillation gain by taking the absolute value (np.abs) of this measure and oscillation phase by taking the angle of this measure (np.angle). To account for phase wrapping, all phase lags above 0 (i.e., phase lead) were subtracted by 2π radians. To estimate the error of gain and phase, we performed a 100-epoch Monte Carlo simulation with replacement at each oscillation frequency for each condition. Error bars derived from all Monte Carlo simulations represent the range of the 95 out of 100 most central epochs relative to the mean value. Shaded error bars unless otherwise noted represent a 95% confidence interval.

---

**Figure S1.**
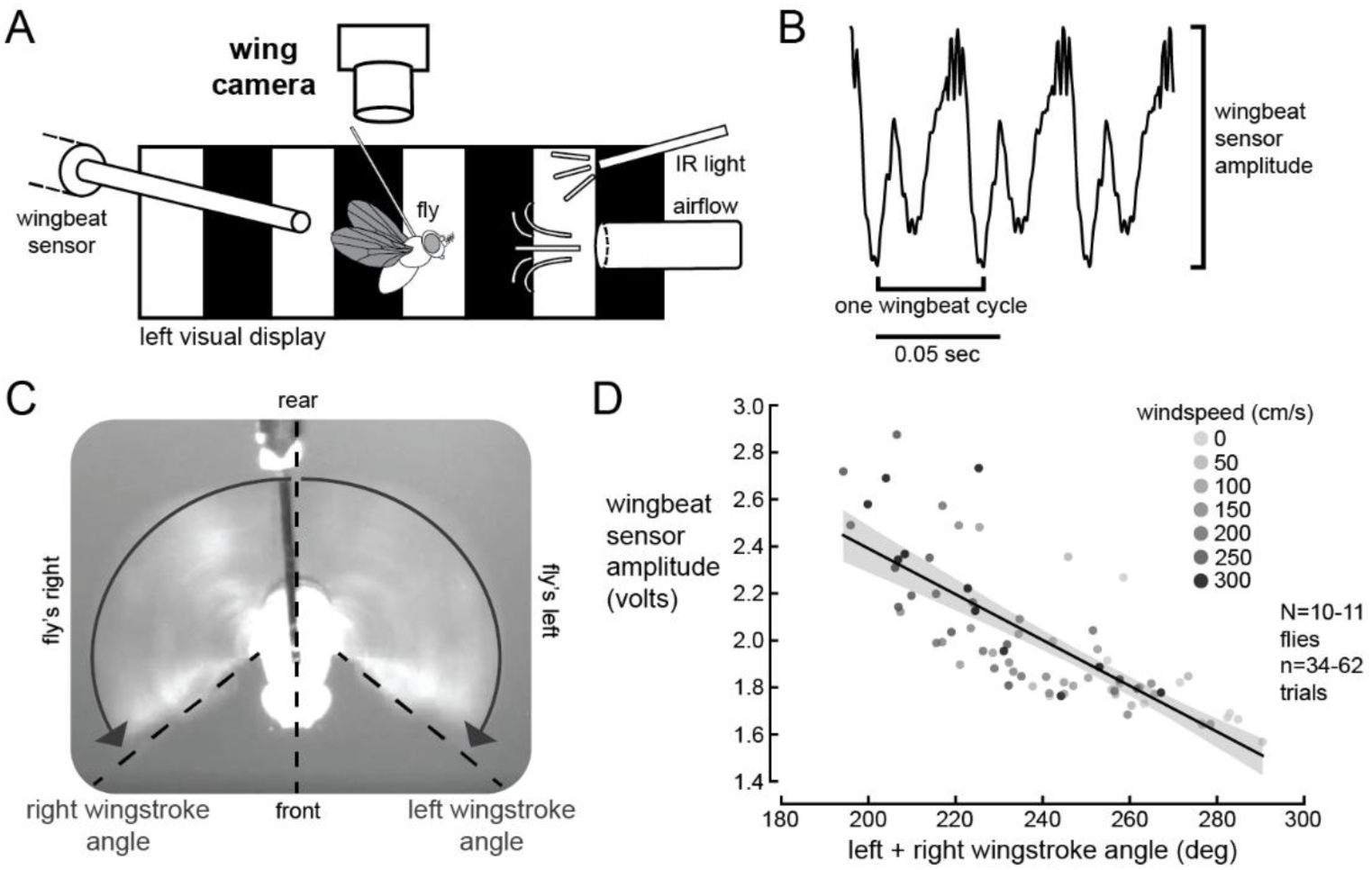
Wingstroke angle is negatively correlated with wingbeat sensor amplitude. A) Schematic of behavioral apparatus (right visual display not shown). Camera capturing wingstroke is directly above the fly, perpendicular to ground. A static grating was present on both the left and right display. B) A short (3 wingbeats) raw wingbeat sensor trace. C) Image from the wingstroke camera. Wingstroke angle is defined as the angle of the leading edge of the wing (dashed lines, diagonal) relative to the midline of the fly (dotted line, vertical). D) Wingstroke angle versus wingbeat sensor amplitude at all wind speeds (N=10-11 flies, n=34-62 trials). This indicates that there is a strong negative, linear correlation between wingstroke angle and wingbeat sensor amplitude (r=-.775, r^2^=.601).

**Figure S2.**
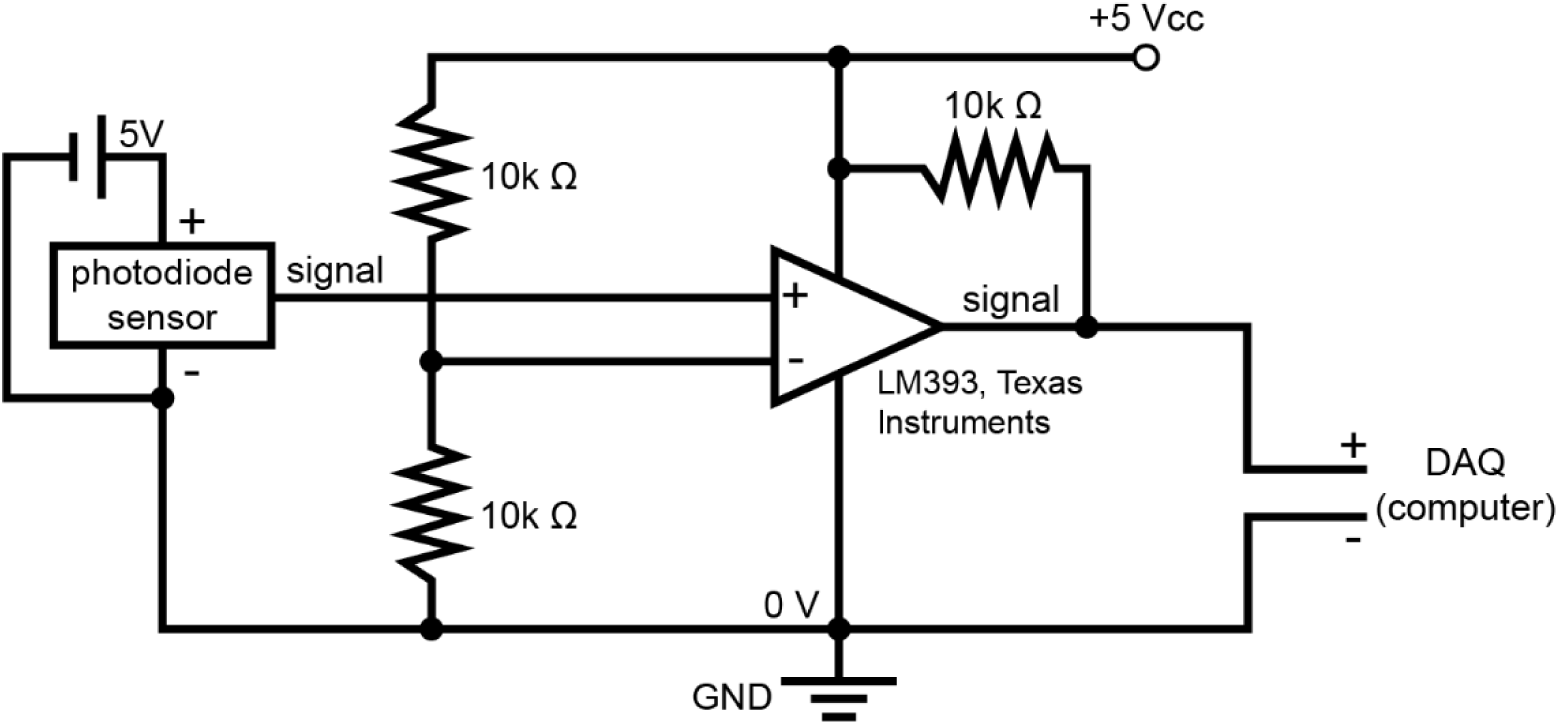
Photodiode-based system for synchronizing wind and visual stimuli. Diagram showing circuit used to synchronize moving static gratings with airflow. A proportional photodiode sensor was used in conjunction with a differential comparator. When the photodiode detected white pixels, a 5 V signal was transmitted to the computer. When the photodiode detected black pixels, no signal was transmitted.

**Figure S3.**
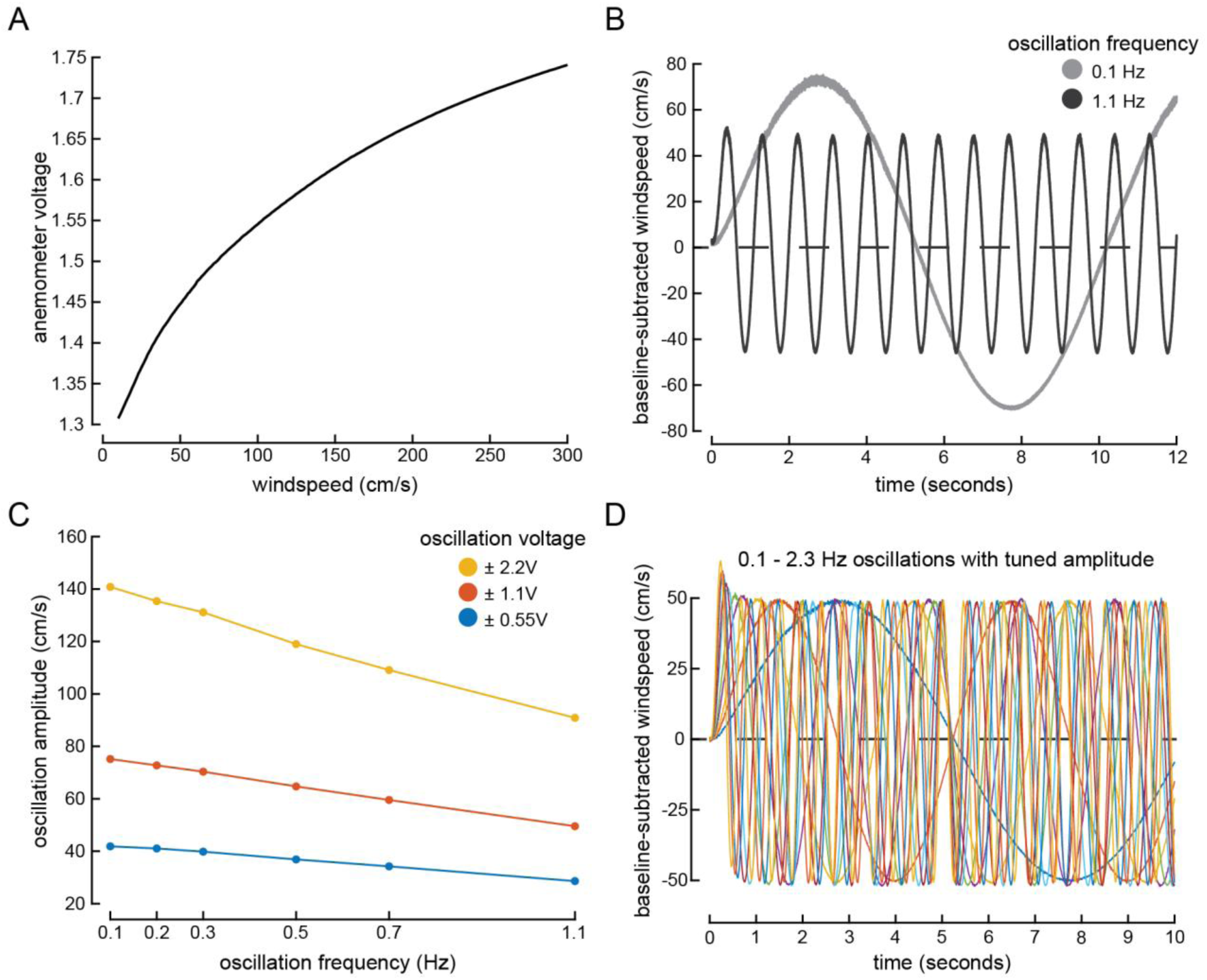
Airflow tuning to maintain wind speed oscillation amplitude across different frequencies. A) Wind speeds presented (in 1 cm/s increments) between 10-300 cm/s and their resulting anemometer values. We presented each wind speed for a 12 sec period, averaging the anemometer value over the last 9 sec. B) Time trace of baseline-subtracted wind speed measurements when the mass flow controller was supplied with ± 1.1 volts at oscillation frequencies of 0.1 Hz (light grey) and 1.1 Hz (dark grey). We determined wind speed values by converting anemometer values using a fitted high-order polynomial curve of (A). C) Amplitudes of wind speed oscillation at frequencies between 0.1-1.1 Hz when the mass flow controller was supplied with ± 2.2 volts (yellow), ± 1.1 volts (red), and ± 0.55 volts (blue). We observed a linearly decreasing relationship at each oscillation voltage for wind speed oscillation amplitude as frequency increased. D) Example post-tuning time traces of baseline-subtracted wind speeds for various oscillation frequencies (0.1, 0.2, 0.3, 0.5, 0.7, 1.1, 1.3, 1.7, 1.9, 2.3 Hz). Across frequencies, the maximum baseline-subtracted wind speed was between 49.70 to 50.96 cm/s and the minimum baseline-subtracted wind speed was between -52.39 to -50.56 cm/s.

**Figure S4.**
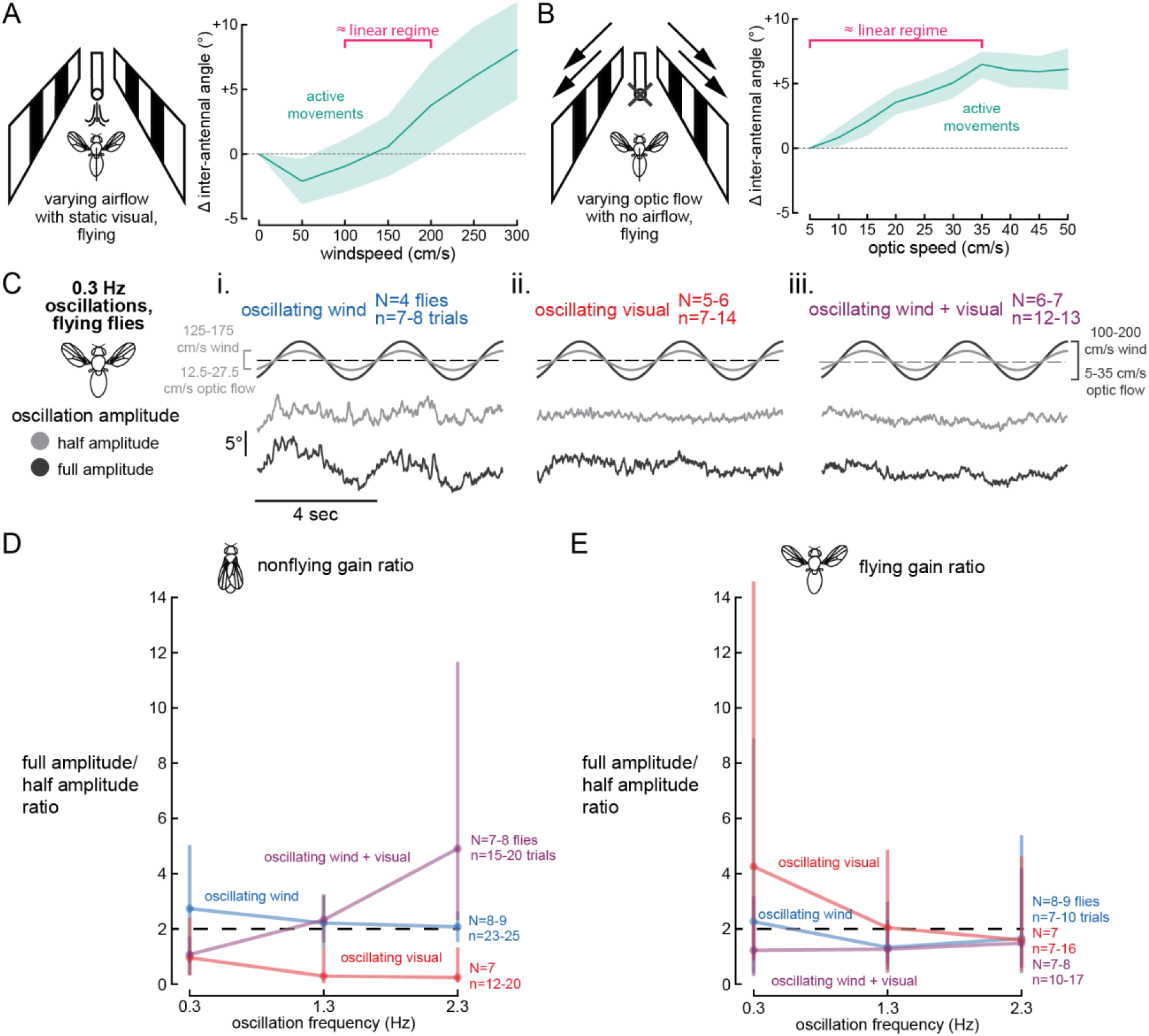
Experimentally measuring linearity and saturation point in the antennal motor system. A) Average active segment inter-antennal angle response to airflow in flying flies presented with a static grating (values are plotted relative to baseline position with no wind). Bracket (pink) highlights the approximately linear wind speed range we used for oscillatory experiments (100-200 cm/s) as it demonstrates a linear pattern across wind speeds. B) Average active segment inter-antennal angle relative to 5 cm/s optic flow in flying flies presented with various optic flow speeds. Bracket (pink) highlights the approximately linear optic flow range we used for oscillatory experiments (5-35 cm/s). C) Average inter-antennal angle response to 0.3 Hz oscillating wind speed (i, with constant 20 cm/s optic flow, blue), 0.3 Hz oscillating optic flow (ii, with constant 150 cm/s airflow, red), and synchronous oscillation of wind speed and optic flow (iii, purple) at full amplitude (dark grey, ± 50 cm/s wind, ± 15 cm/s visual) and half amplitude (light grey, ± 25 cm/s wind, ± 7.5 cm/s visual). D) Ratio of the gain of inter-antennal angle movements with full amplitude stimulus oscillations (dark grey in (C)) to half amplitude stimulus oscillations (light grey in (C)) in nonflying flies with oscillating wind speed (blue), oscillating optic flow (red), and synchronous oscillation of wind speed and optic flow (purple). Linear scaling (doubled antennal amplitude response with doubled oscillation stimulus) would result in a ratio of 2 (dotted line). E) Same as (D), but in flying flies.

**Table S1.** Results of linear-mixed models. Related to Figure 2.

| Figure 2 linear-mixed models |  |  |  |  |
| --- | --- | --- | --- | --- |
| Nonflying inter-antennal angle (2D)<br>iaa ~ windspeed + condition + (1 \| fly) |  |  |  |  |
|  | Coef. (β) | SE | z | p |
| Intercept | -2.34 | 0.853 | -2.74 | 6.16e-3 |
| Condition[static] | 0.923 | 1.20 | 0.767 | 0.443 |
| Windspeed | -0.0159 | 2.61e-3 | -6.09 | 1.10e-9 |
| Windspeed:condition[static] | 0.0478 | 3.66e-3 | 13.1 | 5.96e-39 |
| Group Var | 14.7 | 0.926 |  |  |
| Flying inter-antennal angle (2E)<br>iaa ~ windspeed + condition + (1 \| fly) |  |  |  |  |
|  | Coef. (β) | SE | z | p |
| Intercept | -2.15 | 0.888 | -2.42 | 0.0157 |
| Condition[static] | -0.288 | 1.34 | -0.216 | 0.829 |
| Windspeed | -9.57e-3 | 3.28e-3 | -2.92 | 3.48e-3 |
| Windspeed:condition[static] | 0.0396 | 5.19e-3 | 7.63 | 2.35e-14 |
| Group Var | 12.0 | 0.805 |  |  |
| Wingbeat frequency (2F)<br>wbf ~ windspeed + condition + (1 \| fly) |  |  |  |  |
|  | Coef. (β) | SE | z | p |
| Intercept | 2.06e2 | 2.23 | 92.6 | 0.000 |
| Condition[static] | 7.84 | 3.31 | 2.37 | 0.0180 |
| Windspeed | -0.0254 | 6.23e-3 | -4.08 | 1.21e-4 |
| Windspeed:condition[static] | 0.0372 | 9.69e-3 | 3.84 | 3.53e-5 |
| Group Var | 1.07e2 | 3.16 |  |  |
| Wingbeat amplitude (2G)<br>wba ~ windspeed + condition + (1 \| fly) |  |  |  |  |
|  | Coef. (β) | SE | z | P |
| Intercept | -0.0502 | 0.0510 | -0.985 | 0.324 |
| Condition[static] | 7.59e-3 | 0.0768 | 0.0988 | 0.921 |
| Windspeed | -4.05e-4 | 1.49e-4 | -2.71 | 6.65e-3 |
| Windspeed:condition[static] | -1.79e-3 | 2.37e-4 | -7.56 | 3.99e-14 |
| Group Var | 0.051 | 0.068 |  |  |

**Table S2.**
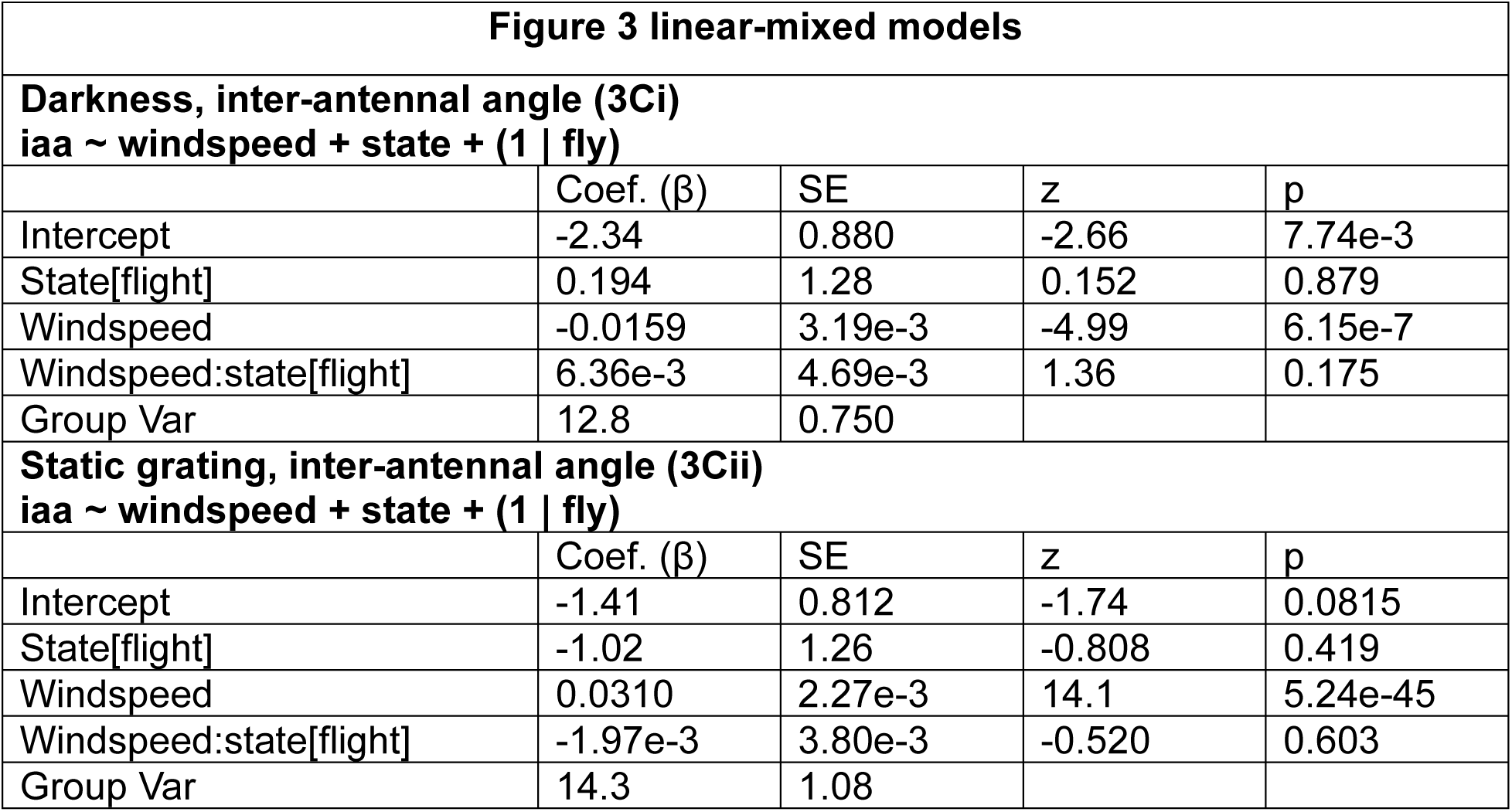
Results of linear-mixed models. Related to Figure 3.

| Figure 3 linear-mixed models |  |  |  |  |
| --- | --- | --- | --- | --- |
| Darkness, inter-antennal angle (3Ci)<br>iaa ~ windspeed + state + (1 \| fly) |  |  |  |  |
| | Coef. ($\beta$) | SE | z | p |
| Intercept | -2.34 | 0.880 | -2.66 | 7.74e-3 |
| State[flight] | 0.194 | 1.28 | 0.152 | 0.879 |
| Windspeed | -0.0159 | 3.19e-3 | -4.99 | 6.15e-7 |
| Windspeed:state[flight] | 6.36e-3 | 4.69e-3 | 1.36 | 0.175 |
| Group Var | 12.8 | 0.750 |  |  |
| Static grating, inter-antennal angle (3Cii)<br>iaa ~ windspeed + state + (1 \| fly) |  |  |  |  |
| | Coef. ($\beta$) | SE | z | p |
| Intercept | -1.41 | 0.812 | -1.74 | 0.0815 |
| State[flight] | -1.02 | 1.26 | -0.808 | 0.419 |
| Windspeed | 0.0310 | 2.27e-3 | 14.1 | 5.24e-45 |
| Windspeed:state[flight] | -1.97e-3 | 3.80e-3 | -0.520 | 0.603 |
| Group Var | 14.3 | 1.08 |  |  |

**Table S3.**
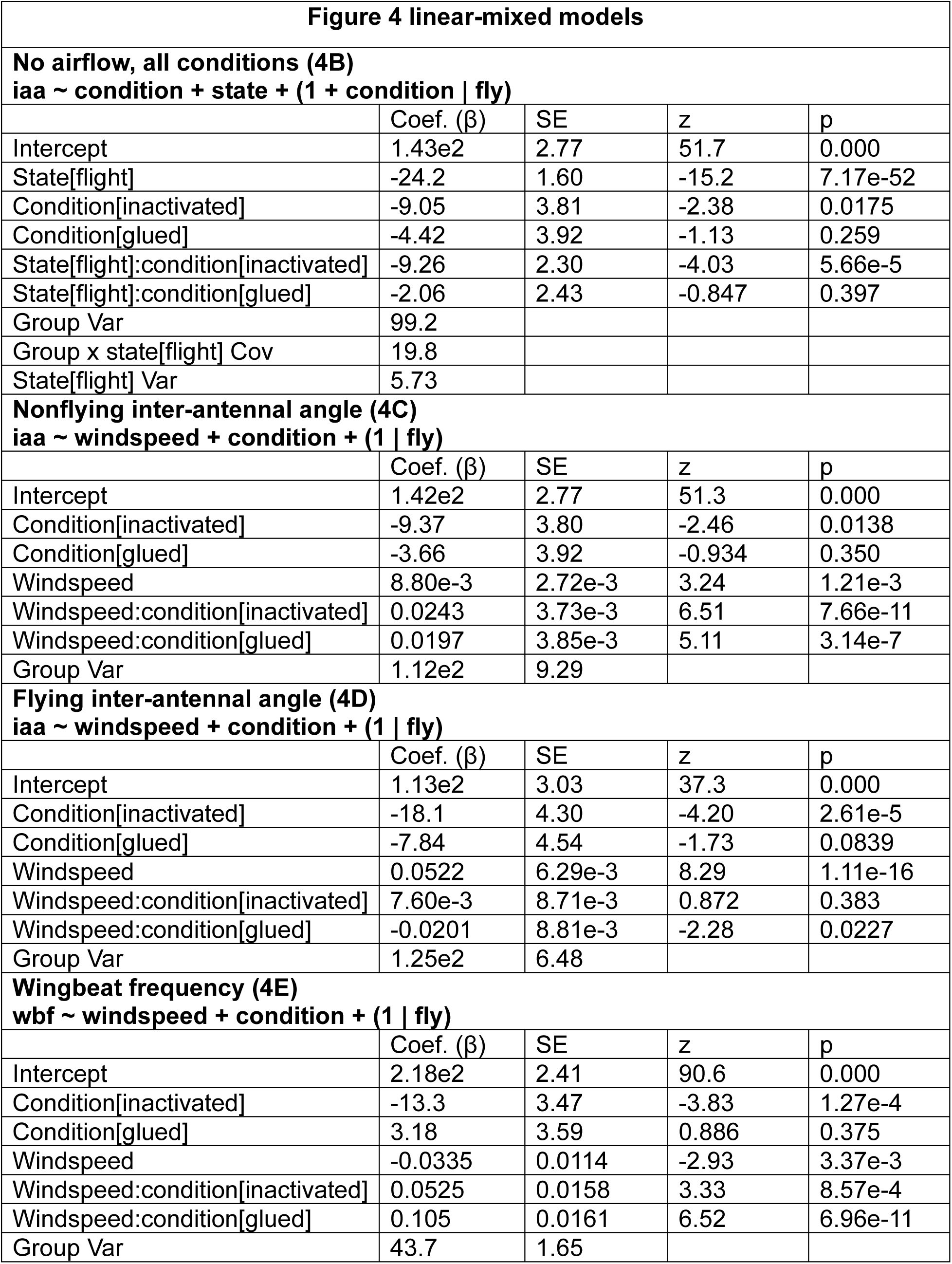
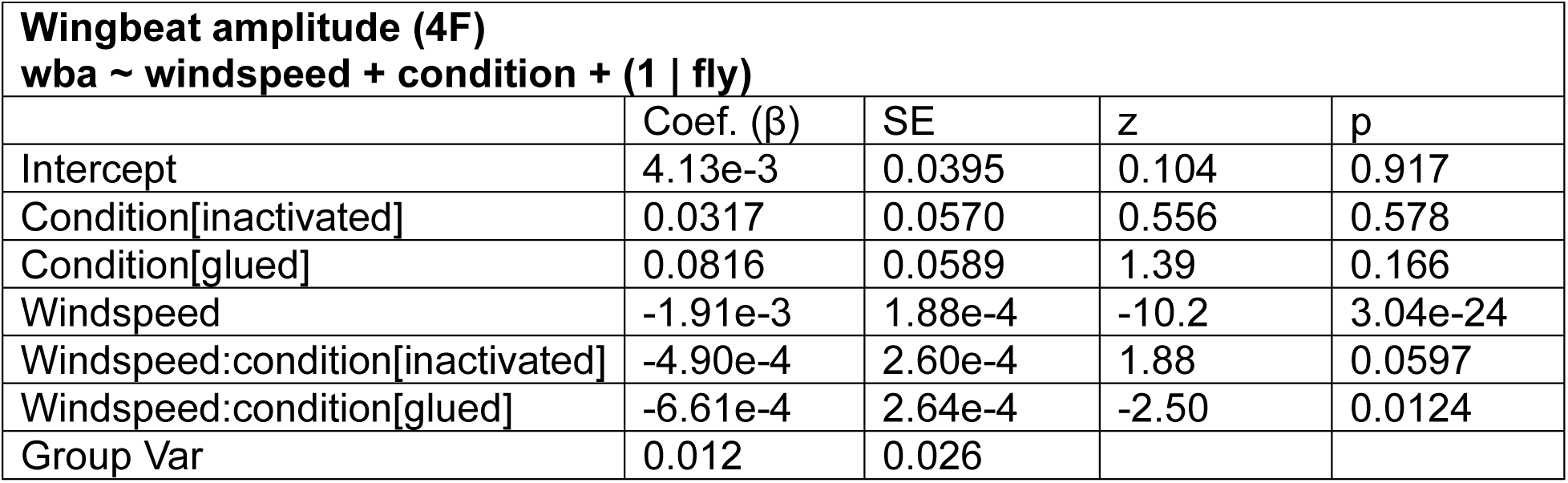
Results of linear-mixed models. Related to Figure 4.

**Figure 4 linear-mixed models**
| **No airflow, all conditions (4B)** |  |  |  |  |
| --- | --- | --- | --- | --- |
| **iaa ~ condition + state + (1 + condition \| fly)** |  |  |  |  |
| | Coef. ($\beta$) | SE | z | p |
| Intercept | 1.43e2 | 2.77 | 51.7 | 0.000 |
| State[flight] | -24.2 | 1.60 | -15.2 | 7.17e-52 |
| Condition[inactivated] | -9.05 | 3.81 | -2.38 | 0.0175 |
| Condition[glued] | -4.42 | 3.92 | -1.13 | 0.259 |
| State[flight]:condition[inactivated] | -9.26 | 2.30 | -4.03 | 5.66e-5 |
| State[flight]:condition[glued] | -2.06 | 2.43 | -0.847 | 0.397 |
| Group Var | 99.2 |  |  |  |
| Group x state[flight] Cov | 19.8 |  |  |  |
| State[flight] Var | 5.73 |  |  |  |
| **Nonflying inter-antennal angle (4C)** |  |  |  |  |
| **iaa ~ windspeed + condition + (1 \| fly)** |  |  |  |  |
| | Coef. ($\beta$) | SE | z | p |
| Intercept | 1.42e2 | 2.77 | 51.3 | 0.000 |
| Condition[inactivated] | -9.37 | 3.80 | -2.46 | 0.0138 |
| Condition[glued] | -3.66 | 3.92 | -0.934 | 0.350 |
| Windspeed | 8.80e-3 | 2.72e-3 | 3.24 | 1.21e-3 |
| Windspeed:condition[inactivated] | 0.0243 | 3.73e-3 | 6.51 | 7.66e-11 |
| Windspeed:condition[glued] | 0.0197 | 3.85e-3 | 5.11 | 3.14e-7 |
| Group Var | 1.12e2 | 9.29 |  |  |
| **Flying inter-antennal angle (4D)** |  |  |  |  |
| **iaa ~ windspeed + condition + (1 \| fly)** |  |  |  |  |
| | Coef. ($\beta$) | SE | z | p |
| Intercept | 1.13e2 | 3.03 | 37.3 | 0.000 |
| Condition[inactivated] | -18.1 | 4.30 | -4.20 | 2.61e-5 |
| Condition[glued] | -7.84 | 4.54 | -1.73 | 0.0839 |
| Windspeed | 0.0522 | 6.29e-3 | 8.29 | 1.11e-16 |
| Windspeed:condition[inactivated] | 7.60e-3 | 8.71e-3 | 0.872 | 0.383 |
| Windspeed:condition[glued] | -0.0201 | 8.81e-3 | -2.28 | 0.0227 |
| Group Var | 1.25e2 | 6.48 |  |  |
| **Wingbeat frequency (4E)** |  |  |  |  |
| **wbf ~ windspeed + condition + (1 \| fly)** |  |  |  |  |
| | Coef. ($\beta$) | SE | z | p |
| Intercept | 2.18e2 | 2.41 | 90.6 | 0.000 |
| Condition[inactivated] | -13.3 | 3.47 | -3.83 | 1.27e-4 |
| Condition[glued] | 3.18 | 3.59 | 0.886 | 0.375 |
| Windspeed | -0.0335 | 0.0114 | -2.93 | 3.37e-3 |
| Windspeed:condition[inactivated] | 0.0525 | 0.0158 | 3.33 | 8.57e-4 |
| Windspeed:condition[glued] | 0.105 | 0.0161 | 6.52 | 6.96e-11 |
| Group Var | 43.7 | 1.65 |  |  |

**Table S3.** Results of linear-mixed models. Related to Figure 4.
| **Wingbeat amplitude (4F)** |  |  |  |  |
| --- | --- | --- | --- | --- |
| **wba ~ windspeed + condition + (1 \| fly)** |  |  |  |  |
| | Coef. ($\beta$) | SE | z | p |
| Intercept | 4.13e-3 | 0.0395 | 0.104 | 0.917 |
| Condition[inactivated] | 0.0317 | 0.0570 | 0.556 | 0.578 |
| Condition[glued] | 0.0816 | 0.0589 | 1.39 | 0.166 |
| Windspeed | -1.91e-3 | 1.88e-4 | -10.2 | 3.04e-24 |
| Windspeed:condition[inactivated] | -4.90e-4 | 2.60e-4 | 1.88 | 0.0597 |
| Windspeed:condition[glued] | -6.61e-4 | 2.64e-4 | -2.50 | 0.0124 |
| Group Var | 0.012 | 0.026 |  |  |

**Table S4.** Results of linear-mixed model. Related to Figure 5.

| Figure 5 linear-mixed model |  |  |  |  |
| --- | --- | --- | --- | --- |
| Inter-antennal angle for nonflying and flying flies (5C-D)<br>iaa ~ optic-speed + state + (1 \| fly) |  |  |  |  |
| | Coef. ($\beta$) | SE | z | p |
| Intercept | 0.166 | 0.357 | 0.465 | 0.642 |
| State[fly] | -0.373 | 0.369 | -1.01 | 0.311 |
| Optic-speed | 0.0196 | 7.39e-3 | 2.65 | 8.11e-3 |
| Optic-speed:state[fly] | 0.123 | 0.0114 | 10.8 | 4.29e-27 |
| Group Var | 2.20 | 0.392 |  |  |

